# High-Throughput Developability Assays Enable Library-Scale Identification of Producible Protein Scaffold Variants

**DOI:** 10.1101/2020.12.14.422755

**Authors:** Alexander W. Golinski, Katelynn M. Mischler, Sidharth Laxminarayan, Nicole Neurock, Matthew Fossing, Hannah Pichman, Stefano Martiniani, Benjamin J. Hackel

## Abstract

Proteins require high developability - quantified by expression, solubility, and stability - for robust utility as therapeutics, diagnostics, and in other biotechnological applications. Measuring traditional developability metrics is low-throughput in nature, often slowing the developmental pipeline. We evaluated the ability of three high-throughput developability assays to predict the bacterial recombinant expression of paratope variants of the protein scaffold Gp2. Enabled by a phenotype/genotype linkage, assay performance for 10^5^ variants was calculated via deep sequencing of populations sorted by proxied developability. We trained a random forest model that predicts expression from assay performance that is 35% closer to the experimental variance and trains 80% more efficiently than a model predicting from sequence information alone. Utilizing the predicted expression, we performed a sitewise analysis and predicted mutations consistent with enhanced developability. The validated assays offer the ability to identify developable proteins at unprecedented scales, reducing the bottleneck of protein commercialization.

## Introduction

A common constraint across diagnostic, therapeutic, and industrial proteins is the ability to manufacture, store, and use intact and active molecules. These protein properties, collectively termed developability, are often associated to quantitative metrics such as recombinant yield, stability (chemical, thermal, and proteolytic), and solubility^1–5^. Despite this universal importance, developability studies are performed late in the commercialization pipeline^2,4^ and limited by traditional experimental capacity^6^. This is problematic because: (i) proteins with poor developability limit practical assay capacity for measuring primary function, (ii) optimal developability is often not observed with proteins originally found in alternative formats (such as display or two-hybrid technologies^7^), and (iii) engineering efforts are limited by the large gap between observation size (~10^2^) and theoretical mutational diversity (~10^20^). Thus, efficient methods to measure developability would alleviate a significant bottleneck in the lead selection process and accelerate protein discovery and engineering.

Prior advances to determine developability have focused on calculating hypothesized proxy metrics from existing sequence and structural data or developing material- and time-efficient experiments. Computational sequence-developability models based on experimental antibody data have predicted post-translational modifications^8,9^, solubility^10,11^, viscosity^12^, and overall developability^13^. Structural approaches have informed stability^14^ and solubility^10,15^. However, many in silico models require an experimentally solved structure or suffer from computational structure prediction inaccuracies^16^. Additionally, limited developability information allows for limited predictive model accuracy^17^. In vitro methods have identified several experimental protocols to mimic practical developability requirements (*e.g*., AC-SINS^18^ and chemical precipitation^19^ as metrics for solubility). However, traditional developability quantification requires significant amounts of purified protein. Noted in both fronts are numerous in silico and/or in vitro metrics to fully quantify developability^1,5^.

We sought a protein variant library that would benefit from isolation of proteins with increased developability and demonstrate the broad applicability of the process. Antibodies and other binding scaffolds, comprising a conserved framework and diversified paratope residues, are effective molecular targeting agents^20–24^. While significant progress has been achieved with regards to identifying paratopes for optimal binding strength and specificity^25,26^, isolating highly developable variants remains plagued. One particular protein scaffold, Gp2, has been evolved into specific binding variants toward multiple targets^27–29^. Continued study improved charge distribution^30^, hydrophobicity^31^, and stability^28^. While these studies have suggested improvements for future framework and paratope residues (including a disulfide stabilized loop), a poor developability distribution is still observed^32^ (Figure 1a,b). Assuming the randomized paratope library will lack similar primary functionality, the Gp2 library will simulate the universal applicability of the proposed high-throughput (HT) developability assays.

**Figure 1.**
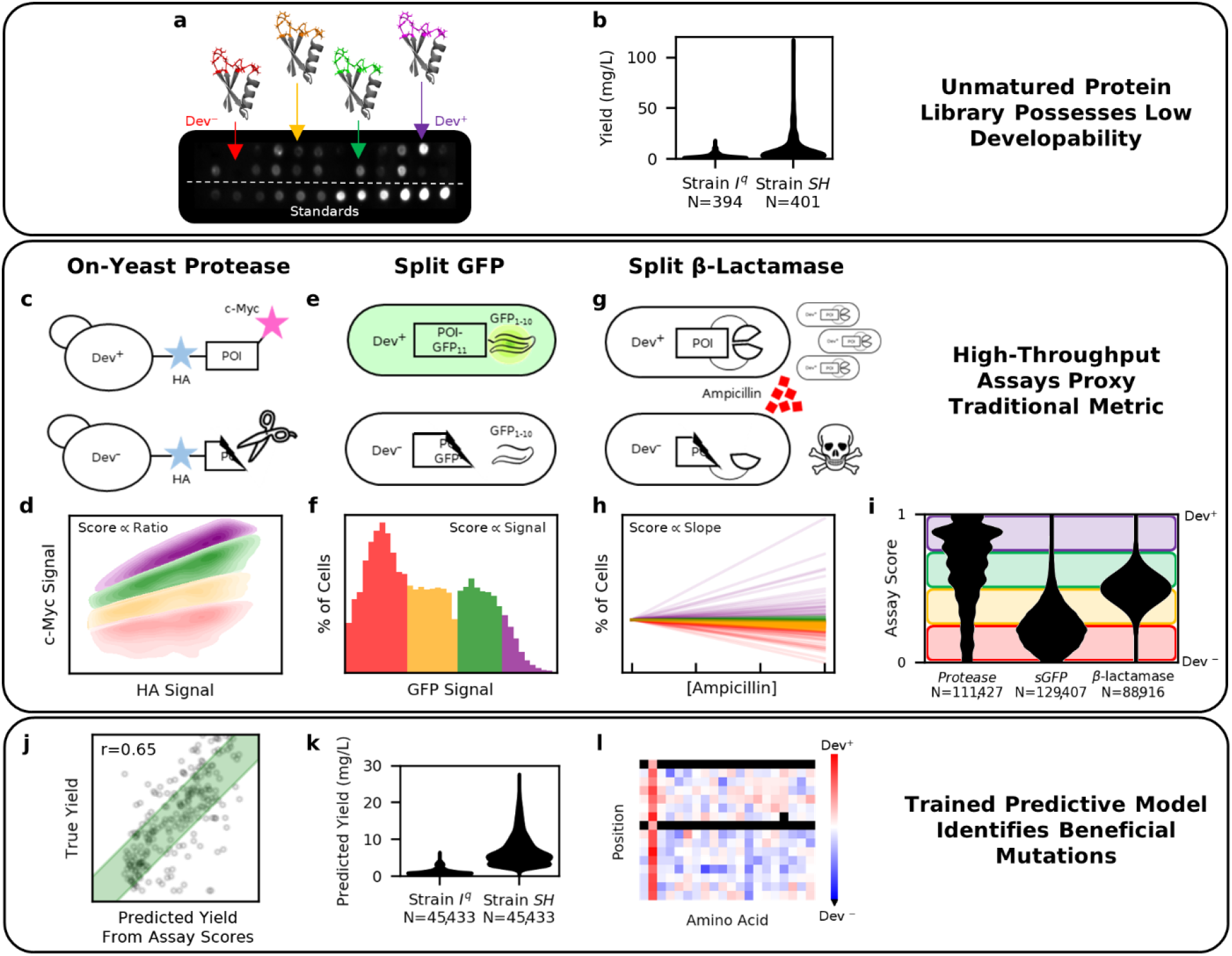
High-throughput (HT) assays were evaluated for the ability to identify protein scaffold variants with increased developability. **a,b)** Gp2 variant expression, commonly measured via low-throughput techniques such as the dot blot shown, highlights the rarity of ideal developability. **c,d)** The HT on-yeast protease assay measures the stability of the protein of interest (POI) by proteolytic extent. **e,f)** The HT split-GFP assay measures POI expression via recombination of a genetically fused GFP fragment. **g,h)** The HT split β-lactamase assay measures the POI stability by observing the change in cell growth rates when grown at various antibiotic concentrations. **i,j)** Assay scores, assigned to each unique sequence via deep-sequencing, were evaluated by predicting expression (see Figure 3). **k,l)** HT assay capacity enables large-scale developability evaluation and can be used to identify beneficial mutations (see Figure 4).

We sought HT assays that allow protein developability differentiation via cellular properties to improve throughput. Variations of three primary assays were examined: 1) On-yeast stability (Figure 1c,d) - previously validated to improve the stability of de novo proteins^33^, antimicrobial lysins^34^, and immune proteins^35^ - measures proteolytic cleavage of the protein of interest (POI) on the yeast cell surface via fluorescence activated cell sorting (FACS). We extend the assay by performing the proteolysis at various denaturing combinations to determine if different stability attributes (thermal, chemical, protease specificity) can be resolved. 2) Split green fluorescent protein (GFP, Figure 1e,f) - previously used to determine soluble protein concentrations^36^ - measures the assembled GFP fluorescence emerging from a 16-amino acid fragment (GFP_11_) fused to the POI after recombining with the separably expressed GFP_1-10_. We extend the assay by utilizing FACS to separate cells with differential POI expression to increase throughput over the plate-based assay. 3) Split β-lactamase (Figure 1g,h) - previously used to improve thermodynamic stability^37^ - measures cell growth inhibition via ampicillin to determine functional lactamase activity achieved from reconstitution of two enzyme fragments flanking the POI. We expand assay capacity by deep sequencing populations grown at various antibiotic concentrations to relate change in cell frequency to functional enzyme concentration.

In this paper, we determined the HT assays’ abilities to predict Gp2 variant developability. We deep-sequenced the stratified populations and calculated assay scores (correlating to hypothesized developability) for ~10^5^ Gp2 variants (Figure 1i). We then converted the assay scores into a traditional developability metric by building a model that predicts recombinant yield (Figure 1j). The assays’ capacity enabled yield evaluations for >100-fold traditional assay capacity (Figure 1k, compared to Figure 1b) and facilitated protein engineering by observing beneficial mutations in highly developable proteins (Figure 1l).

## Results

### Gp2 Paratope Library Quantification

We first evaluated the assays’ ability to separate sequence classes with a hypothesized difference in developability. 204,174 observed Gp2 variants belonged to one of four classes (Figure 2a): *GaR*: a thermostable variant^27^, *Stop*: 13,690 nonfunctional truncated variants; *CC+*: 128,854 variants with a hypothesized^28^ stabilizing cysteine pair at sites 7 and 12; *CC−*: 61,629 variants without conserving sites 7 and 12. *CC+, CC−*, and *Stop* classes utilize a previously optimized conserved framework^28^ and two paratope loops, each with 6-8 ‘NNK’ degenerate codons encoding all 20 amino acids (Figure 2b). The library was widely diversified, averaging 13.1 differences between observed sequence pairs (Figure 2c).

**Figure 2.**
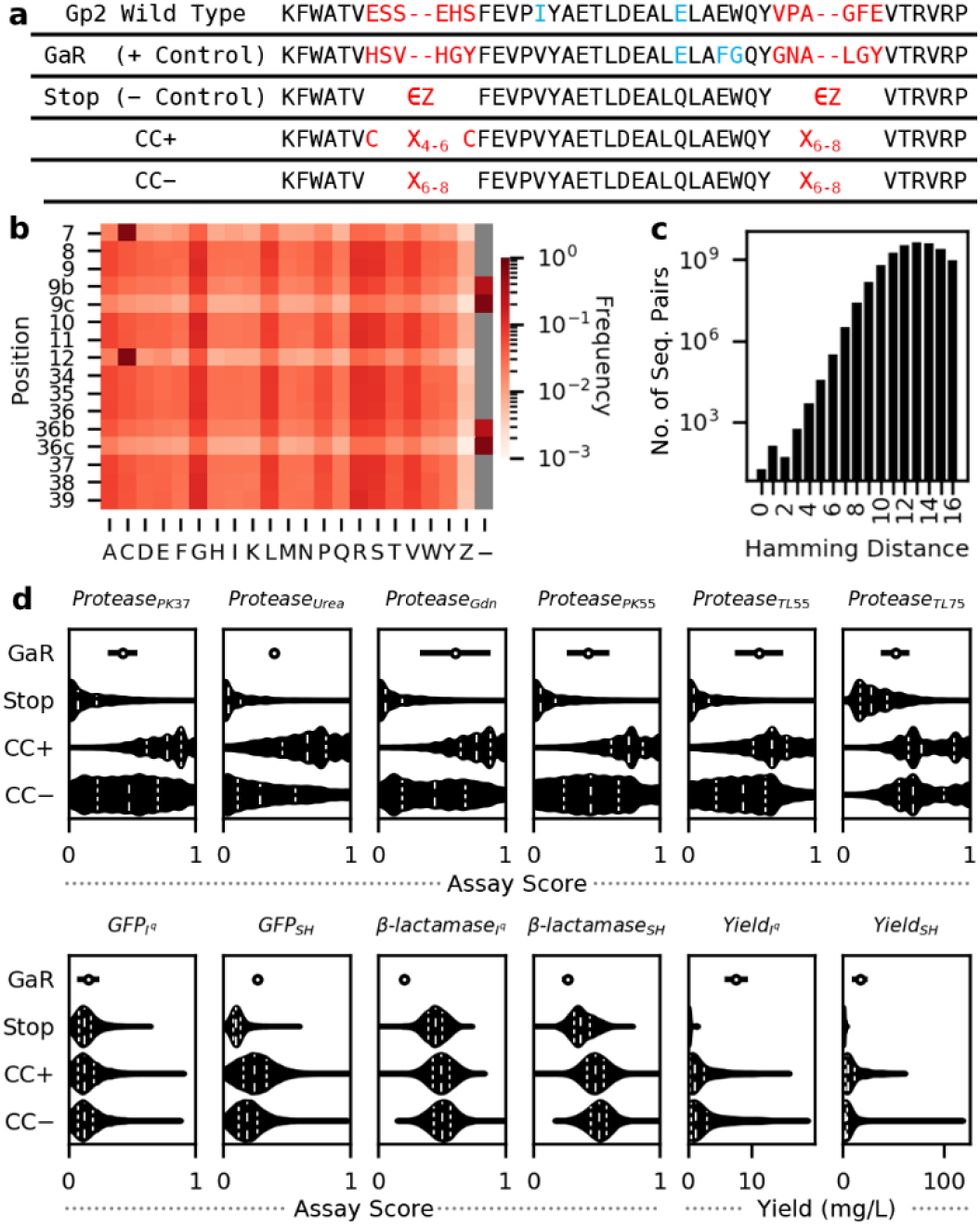
Developability Characterization of Loop-Diversified Gp2 Library. **a**) Sequence alignment of assayed sequence classes: *GaR* (single variant control), *Stop*: (sequences with stop codon, Z), *CC+*: (hypothesized to be more developable), *CC−*: (hypothesized to be less developable) **b**) Diversified paratope frequency heatmap. **c**) Histogram depicting the pairwise distances between 190,483 full length and unique variants. **d)** Assay performance distributions divided by class. *Top Row*: various on-yeast protease assay reaction conditions. *Bottom Row*: bacterial assays performed in strain I^q^ and strain SH. GaR error bars represent the standard deviation (n=3 trials). Total unique variants for Stop, CC+, and CC− range 93,178-140,229 for HT assays, and 431-447 for yield (See Supplementary Figure 2)

### Recombinant Yield as a Traditional Developability Metric

We sought a traditional developability metric that was translationally relevant and scalable to train and validate predictive models. A key step in developing and using a protein involves recombinant production. Bacterial cells are often chosen due to affordability, ease, and speed^38^. However, with limited production machinery, expressed proteins must rely on inherent developability parameters to achieve high soluble concentrations. Also, considering alternative assays require high purified protein quantities, we selected bacterial recombinant yield as the metric of interest. The Gp2 titer in the soluble lysate fraction was measured using a chemiluminescent quantitative dot blot protocol^39^ via a C-terminal His_6_ tag (Figure 1a,b).

Different bacterial strains have been evolved containing additional machinery to obtain increased yield. We chose to include two *E. coli* strains (T7 Express lysY/I^q^ (I^q^) and SHuffle^®^ T7 Express lysY (SH), New England Biolabs) for improved developability resolution. SH was chosen to stabilize disulfide formation and increase cysteine-free variant yields^40^. This was confirmed by *GaR* having a significantly higher yield in SH despite not having cysteines (p<0.05 in one-way Student’s t-test using trial-averaged yield, n=8 plates per strain).

The recombinant yield of unique Gp2 sequences in each class was measured in triplicate (Figure 2d): *GaR* (both strains), *Stop* (I^q^: 37 Gp2 variants, SH: 46), *CC−* (I^q^: 98, SH: 117), and *CC+* 296, SH: 284). *GaR* had a significantly higher yield than most *Stop* sequences (I^q^: 100%, SH: 63% of unique *Stop* sequences, p<0.05 in one-way Student’s t-test using plate-averaged *GaR* standard deviation, n=3 trials), validating the dot blot controls while suggesting slight noise with SH. *CC+* did not have significantly different yields than *CC−* (p = 0.40 in two-way Mann-Whitney U test) in I^q^, while the populations were significantly different in SH (p<0.05, one-way Mann-Whitney U test). This implies SH is forming a disulfide bond, thus increasing *CC+* sequence developability.

### HT Developability Assays

The Gp2 variants were sorted into populations of varying developability and were assigned an HT assay score as the mean over three independent trials (Supplementary Figure 1). Below we motivate score calculation, followed by assay score distribution analysis (Figure 2d, Supplementary Figure 2).

#### On-yeast Stability

The on-yeast stability assay evaluates protein stability by measuring proteolytic cleavage (Figure 1c). Using yeast surface display technology^41^, the POI is expressed between two tags (N-terminal HA, C-terminal cMyc). The protein-displaying yeast are exposed to a protease at a concentration that produces a distribution of cleavage (as determined by cMyc:HA ratio) across protein variants. The Gp2 library was sorted into four populations (Figure 1d). Sequencing scored every collected variant on a cell-weighted average: 1 (intact), 2/3, 1/3, 0 (fully cleaved).

We performed the proteolysis using various conditions to determine if additional stability metrics could be obtained (Supplemental Figure 1). From our baseline condition (P_PK37_), we studied chemical stability by adding 1.5 M urea (P_Urea_) or 0.5 M guanidinium chloride (P_Gdn_). We explored protease specificity by using proteinase K (P_PK55_) and thermolysin (P_TL55_). Finally, we examined thermostability for each enzyme at an additional temperature (P_PK37_ vs. P_PK55_ and P_TL55_ vs. P_TL75_).

Assay scores were calculated for >10^5^ unique Gp2 variants in each of the 6 reaction conditions. The assay score distributions per class (Figure 2d) matched hypothesized developability in all conditions except P_TL75_. Standard deviations were small (0.17 – 0.20, except P_TL75_: 0.29). *Stop* variants scored low (0.04 – 0.08, except P_TL75_: 0.23). *GaR* scored higher than most *Stop* variants (67 – 81%, except P_TL75_: 35%). One potential hypothesis for P_TL75_ is the increased temperature may lead to non-specific binding of surface-aggregated proteins. Nevertheless, all reaction conditions, displayed a significantly higher distribution of assay scores for *CC+* vs. *CC−* (one-way Mann-Whitney U test, p<0.001), validating each condition’s utility.

#### Split GFP

The split GFP assay measures POI concentration with a C-terminus fused 11^th^ strand of GFP (Figure 1e). Upon recombination with GFP strands 1-10, which was separately induced following POI production and a one-hour gap, the POI fusion remaining soluble in the cytosol will produce a fluorescent signal detectable by FACS (Figure 1f). The library was sorted into four populations based on GFP signal and assigned an assay score as a cell-weighted average: 1 (highest signal), 2/3, 1/3, 0 (background signal).

The assay score distributions (Figure 2d) are consistent with expectations in SH (G_SH_) with limited resolution in I^q^ (G_Iq_). While both distributions display a low assay score skew, *GaR* had a significantly higher score than 76% of *Stop* in G_SH_, compared to 8% in G_Iq_. Additionally, G_SH_ produced a significantly higher assay score distribution for *CC+* compared to *CC−* (one-way Mann-Whitney U test. p<0.001) whereas G_Iq_ scores were only nominally higher (p=0.15). Thus, G_SH_ is a compelling candidate for HT developability analysis.

#### Split β-lactamase

In the split β-lactamase assay, the POI is inserted in a loop distal to the active site (final construct: β-lac_1-194_-(G_4_S)_2_-AS-POI-GS-(G_4_S)_2_-β-lac_197-287_, location previously observed to retain 40% activity^42^). Functional enzyme, hypothesized to be paired with POI solubility and folding robustness^43^, provides ampicillin resistance allowing cell reproduction (Figure 1g). The change in growth rates was measured as the change in POI amplicon abundance in cultures grown to saturation with varying antibiotic concentrations (Figure 1h). For comparison to other assays and improved modeling efficiency, slopes were normalized and scaled (see Methods).

The split β-lactamase assay produced assay scores that were contradictory towards hypothesized developability yet were able to differentiate classes, suggesting potential utility despite an unsolved mechanism. We obtained assay scores for 1×10^5^ variants in both I^q^ (β_Iq_) and SH (β_SH_). Independent *GaR* cultures (capable of growing at all concentrations) and *Stop* (unable to grow in non-zero ampicillin concentrations) performed as expected (Supplementary Figure 3). Yet in multi-POI culture, *GaR* had a significantly lower assay score than *Stop* (β_Iq_: 99%, β_SH_: 70%, one-way Student’s t-test, p<0.05), and the *CC+* population had a significantly lower assay score distribution than *CC−* (both strains, one-way Mann-Whitney U test, p<0.001). See Figure 5 and Discussion for further explanation.

### Determination of Most Predictive HT Assay Conditions

While the HT assays broadly differentiated hypothesized class developability, the ability to transform the assay scores to a traditional metric is a superior utility assessment. We observe that despite the limited sensitivity in the split GFP assay and the counterintuitive split β-lactamase distributions, the assays have nonzero mutual information (MI) with yield, suggesting utility as long as the predictive model is capable of exploiting the nonlinear relationships captured by MI (Supplementary Figure 4, see Figure 5c). In this section, we determine the optimal HT assay set (assay type, reaction conditions, and/or bacterial strain) by the ability to predict recombinant yield with the lowest mean-squared error (MSE) loss.

With a potential complex relationship between developability and assay scores, we designed our model to maximize the ability to detect assay utility. Correlation of yields in both strains was observed (ρ *CC+*: 0.65, *CC−*: 0.61; Supplementary Figure 5); thus, a multitask model (Figure 3a) was utilized to include both strains’ yield measurements via a one-hot (OH) encoded vector. We included relevant comparisons for model inputs: a null strain-only model (predicts the mean yield per strain) and a OH sequence model (encoded and flattened paratope sequence). To capture possible linear and nonlinear relationships between assay scores, sequences, strains, and yield, four model architectures (ridge, random forest, support vector machine, and a feedforward neural network) were employed.

**Figure 3.**
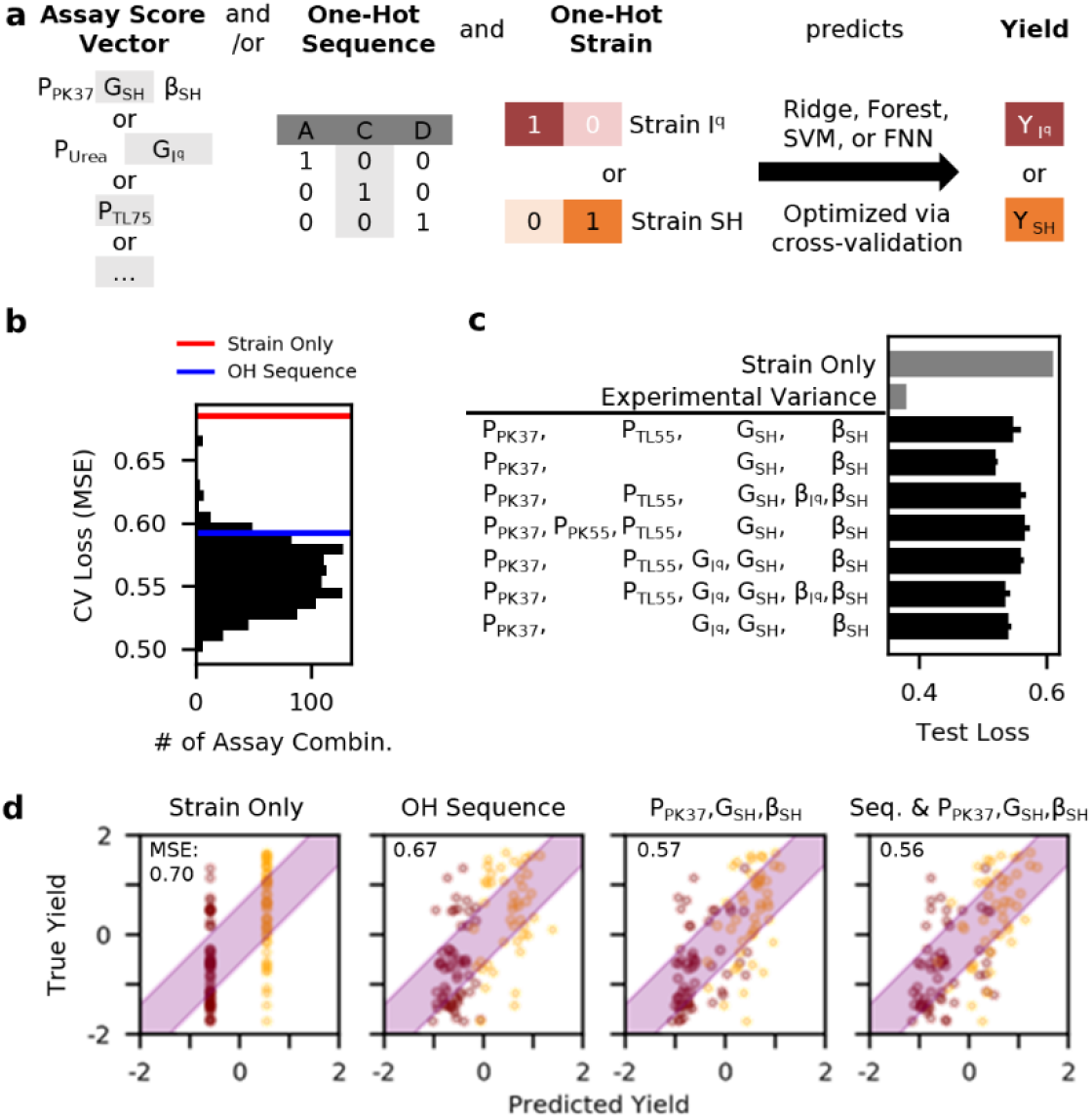
Determination of Predictive HT Developability Assays. **a**) Model visualization utilizing HT assay scores, a one-hot paratope sequence, and a one-hot strain identifier to predict the recombinant yield in both cell types. **b**) Predictive model loss (mean squared error (MSE) between predicted and actual yields) distribution for 1,023 HT assay combinations. **c)** The top combinations from CV (listed top down) were tested for generalizability by the predictive loss against independent set of 44 sequences. **d)** Representative scatter plots of predicted versus measured yield (I^q^: purple, SH: orange) during final evaluation on set of 97 sequences. Purple shaded area represents true yield ± square root of sequence-averaged experimental variance.

Cross-validation (CV) and hyper-parameter optimization were trained by 195 unique sequences observed in all HT assays and for which yield was measured in at least one strain. A Yeo-Johnson^44^ power transform and normalization was applied to remove correlation between error and yield (λ = −0.324, Supplementary Figure 6). The experimental variance (measurement accuracy) was calculated as the sequence-averaged trial-to-trial (n=3) variance after applying the transformation to trial yields.

Despite potential limitations, all 1023 assay combinations of the 10 HT conditions predicted yield with a lower CV loss than the strain-only control, and 92% of the combinations outperformed the OH sequence model (Figure 3b) suggesting all conditions possess utility. There were seven assay combinations (using seven of the ten assays) that performed optimally and equally (Supplementary Figure 6, one-way Student’s t-test against top model, p>0.05). To determine the most generalizable collection, the yield for an independent set of 44 sequences (not utilized during CV but observed in top seven HT assays) was predicted revealing the most informative set: P_PK37_, G_SH_, β_SH_ (Figure 3c, one-way Student’s t-test against top model, p<0.05).

The top three HT assays can provide substantial predictive power for variant developability over sequence information alone. The yield for a second set of 97 sequences (not utilized during CV but observed in top three HT assays) was predicted (Figure 3d, Supplementary Figure 7). The assay model (MSE: 0.565) was able to significantly (one-way Student’s t-test, p<0.05) outperform the one-hot sequence model (MSE: 0.667) and strain-only model (MSE: 0.697). A model utilizing both sequence and assay information (MSE: 0.562) did not have significantly different (p>0.05) performance from the assay model alone, suggesting little aid of sequence knowledge as currently implemented. The model utilizing sequence and assay information, while predicting better than alternatives, required a nonlinear random forest architecture with 325 trees for optimal predictive performance that still trails the experimental variance (0.364), suggesting room for future improvement. As performed, the assays reduce the gap between prediction and experimental error of developability evaluation by 35% compared to sequence information alone.

### Optimal Paratope Sequence Identification

With a predictive model to translate the assay scores to recombinant expression, we aimed to understand the sequence-developability relationship. The predictive model utilizing P_PK37_, G_SH_, β_SH_ assay scores and OH sequence was used to predict the yield for 45,433 unique sequences in both strains (Figures 1k & 4a). After observing the predicted yield distribution, 6,394 sequences with a predicted I^q^ yield > 2.5 mg/L (transformed yield > 0.0) and SH yield > 6.4 mg/L (transformed yield > 0.75) were isolated as Dev^+^. The pairwise Hamming distance distribution for the Dev^+^ sequences (median 12.3) compared to the initial distribution (median 13.0) significantly (χ^2^, p < 0.05), suggesting that developable sequences exist in a partially constrained subset of sequence space.

**Figure 4.**
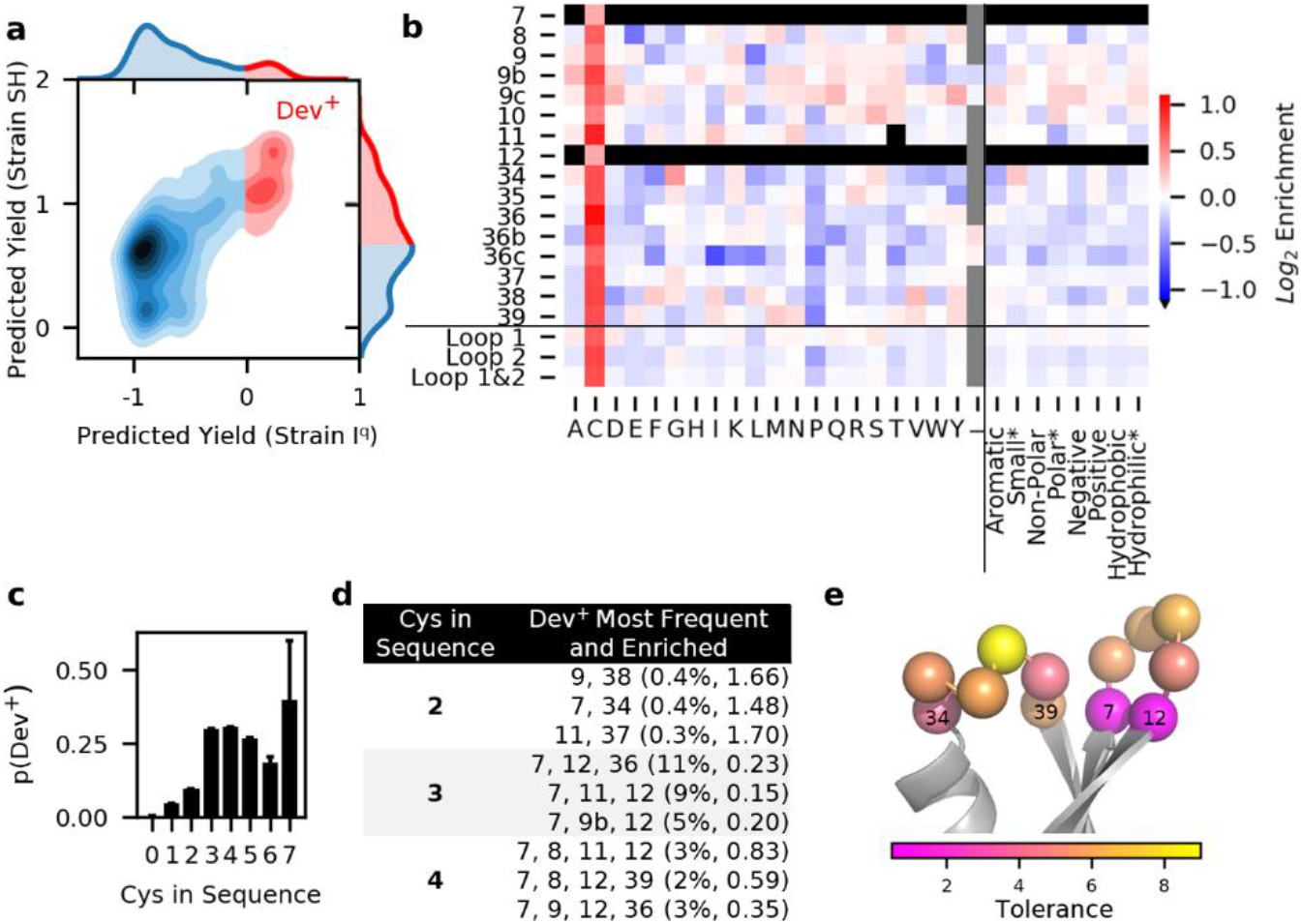
HT Assays Enable Prediction of Gp2 Variants with High Developability. **a**) Kernel density plot of the predicted yield of 45,433 unique sequences in each bacterial strain. 6,394 sequences with high predicted yield in both strains were isolated as Dev^+^ (red). **b)** Sitewise enrichment heatmap (Dev^+^ versus all predicted sequences) for each amino acid and averaged groups with similar chemical properties: aromatic (F, W, Y), small* (A, G, S), non-polar aliphatic (A, G, I, L, M, P, V), polar uncharged* (N, Q, S, T), negative charged (D, E), positive charged (H, K, R), hydrophobic (A, F, G, I, L, M, P, V, W, Y), and hydrophilic* (D, E, H, K, N, Q, R, S, T). *Note: cysteine was removed to identify any further enrichment of the groups. Loop 1: positions 8-11. Loop 2: positions 34-39. **c)** The proportion of sequences predicted identified as Dev^+^ as a function of the number of cysteines in the sequence. Error bars: 1 / number of predicted sequences. **d)** The most frequent (percent of Dev^+^) and enriched (log2 of Dev^+^ versus all predicted) positions for combinations of cysteines that result in high developability proteins. **e)** Wild type paratope positions of Gp2 (PDB: 2WMN) colored by the mutational tolerance calculated as the inverse of the average magnitude of amino acid enrichment.

To identify beneficial, tolerable, and detrimental mutations to developability, the log2 difference in amino acid frequency at each position between Dev^+^ and all predicted sequences was calculated (Figure 4b). Cysteine was the only positively enriched amino acid at positions 7 and 12 (confirming *CC+* stability) but was also the most enriched at every position. The high cysteine enrichment was also observed when analyzing predictions of an assay score model without sequence information (Supplementary Figure 8). Regarding epistasis, we analyzed the probably of Dev^+^ as conditioned by number of cysteines in the sequence, finding 3 cysteines most optimal (Figure 4c). To determine the best cysteine locations to improve developability, the Dev^+^ frequency and log2 enrichment were calculated (Figure 4d). It should be noted the 7 and 12 pair had a negative enrichment likely due to the artificially increased initial frequency. As additional cysteines may be disfavored for downstream processing flexibility, the enrichment of sequences only containing cysteines at positions 7 and 12 was calculated (Supplementary Figure 9). Enabled by the assay throughput, less-extreme enrichment values observed for cysteine-rich sequences (compared to sequences with fewer cysteines) suggests the cysteines are buffering stability and permitting a wider sequence set.

Additional analysis enabled by the HT assays were used to hypothesize properties that drive Gp2 stability. The enrichment of small residues (alanine, glycine, and serine) at position 34, the proline depletion in the second loop, and gap enrichment at positions 36b,c (enriching sequences of wild type length) suggests the second loop may be geometrically constrained. We assessed positional mutational tolerance (ability to mutate without modifying developability) by calculating the inverse of the average enrichment score magnitude (Figure 4e). Positions 7 and 12 were the most constrained (tolerances: 0.5), signifying the need to be cysteines. While position 37 was the least constrained position (8.8), as a whole loop 2 (5.5) was less tolerant than loop 1 (5.9, excluding 7 and 12). We hypothesize either i) the second loop is a poor paratope in terms of allowing broad diversity with favorable developability or ii) the stabilizing disulfide bond offsets unfavorable mutations within the first loop.

### ϐ_SH_ Assay Predictive Performance Explained by Mutual Information

Like amino acid preference, we sought a first-order understanding of optimal assay scores by looking at the Dev^+^ distribution compared to all observed unique sequences (Figure 5a). Matching the sequence class distributions (see Figure 2), P_PK37_ and G_SH_ assay scores of Dev^+^ sequences were significantly higher, and β_SH_ assay scores was significantly lower than the initial distribution (Figure 5a, one-way Mann-Whitney U test, p<0.05). However, the rank correlation between β_SH_ and yield is slightly positive (Iq: 0.00, SH: 0.11), suggesting the model is exploiting a nonlinear relationship.

**Figure 5.**
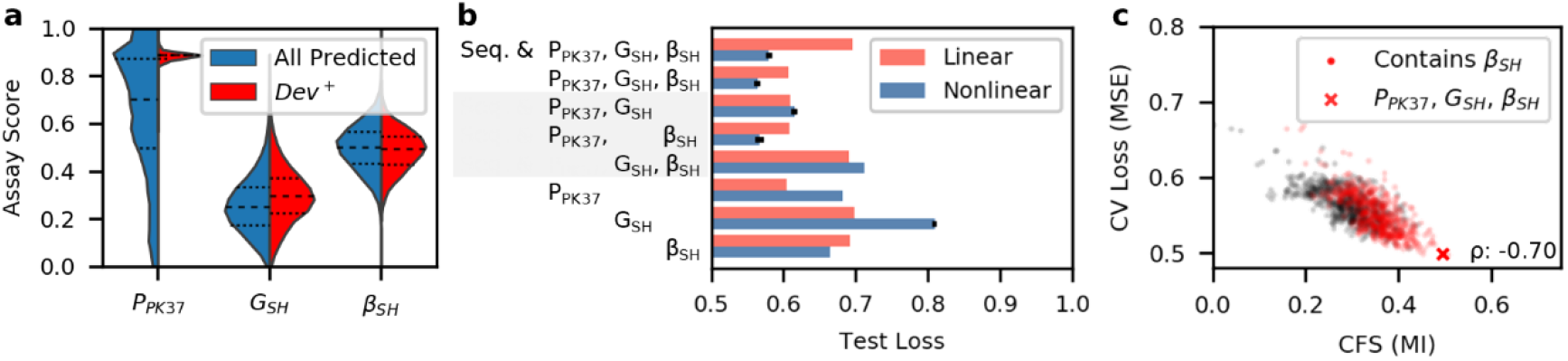
Nonlinear models can extract nonlinear developability mutual information (MI) from the split β-lactamase assay. **a**) Comparison of assay score distributions between 45,433 unique sequences with observed P_PK37_, G_SH_, β_SH_ assay scores (blue) versus 6,394 of the sequences with high predicted developability (Dev^+^, red). **b)** The predictive performance of model input combinations in both a linear architecture (ridge regression) and nonlinear architectures (reported top performance of random forest, support vector machine, and a feed-forward Neural Network). Error bars in nonlinear models represent standard deviation in MSE from n=10 stochastically trained models. **c)** The correlation-based feature selection (CFS) as calculated by MI for 1023 assay combinations versus the CV loss utilizing the best of linear and nonlinear model architectures. The Spearman’s rank correlation coefficient (ρ) between CFS and loss confirms the ability of the models to extract nonlinear MI.

We hypothesize that the counterintuitive relationships between β_SH_ and yield resulted from several competing interactions relating the change in sequence frequency to the concentration of functional enzyme-POI. We tested this by comparing nonlinear versus linear model performances for several model input combinations (Figure 5b). While the P_PK37_ and G_SH_ assays, alone and together, performed better with a linear model, 4 of 5 models using the β_SH_ assay performed best with a nonlinear model.

The correlation-based feature selection^45^ (CFS) explains how the nonzero MI between β_SH_ and yield (I^q^: 0.16, SH: 0.13) resulted in increased predictive power by supplying non-redundant information with respect to other HT assays. The CFS calculated by MI was significantly higher and CV loss was significantly lower of HT assay combinations containing β_SH_ than assay combinations without (Figure 5c, one-way Mann-Whitney U test, p<0.05). CFS calculated with MI was highly correlated with loss when utilizing nonlinear models (ρ = −0.70) remarking its effectiveness as a feature selection tool. We also found CFS calculated by rank correlation was correlated to linear model performance (ρ = −0.56) but less so to overall performance (ρ = −0.30) as linear models cannot exploit nonlinear relationships (Supplementary Figure 4). As a result, the top CFS combination via rank correlation (P_PK37_, P_Urea_, P_PK55_, G_Iq_, G_SH_. Ridge MSE: 0.564) increased the prediction error relative to experimental variance by 46% compared to the top model identified by CFS via MI (P_PK37_, P_TL55_, G_SH_, β_SH_. Forest MSE: 0.497).

### Training Sample Size Evaluation

Finally, we asked how the predictive performance scales versus the number of training sequences. We first analyzed how many sequences it takes for a model to learn training set developability, as determined by outperforming the strain only model during CV (Figure 6a). With only 10 sequences (5% of data), the P_PK37_, G_SH_, β_SH_ model achieves this goal (one-way paired t-test, p<0.05). However, models with sequence information required at least 39 sequences (20% of data) to achieve the same accomplishment, suggesting the increased input dimensionality limits the model’s ability to learn. When evaluating the models for generalizability against a test set (Figure 6b), the models using assays required only 59 (P_PK37_, G_SH_, β_SH_, 30% of data, p<0.05) or 78 (Sequence and P_PK37_, G_SH_, β_SH_, 40% of data, p<0.05) training sequences to outperform the strain only model, while the sequence only model required all 195 sequences. The generalizability results suggest the HT assays reduce the training data requirements by 60-70% over sequence information alone.

**Figure 6.**
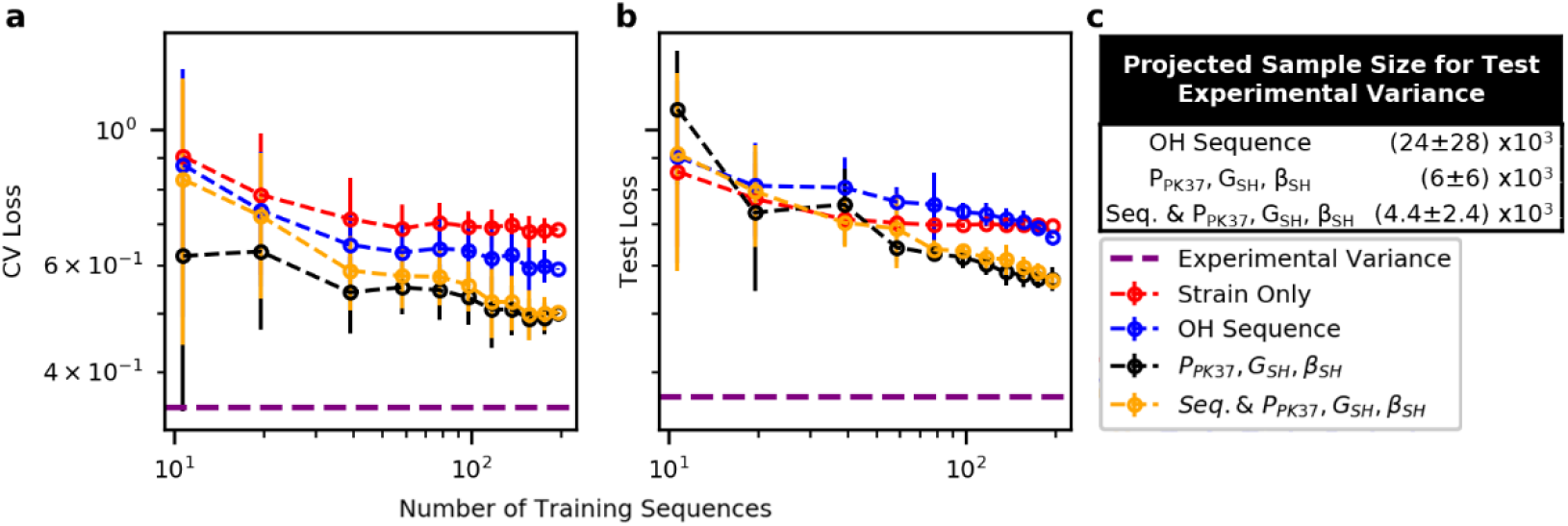
HT developability assays reduce training size requirement. Ten bootstrapped samples for each sample size (ranging 5-100% of available data) were individually trained by CV and evaluated on 97 independent test sequences. Error bars represent standard deviation across models. **a**) The performance during CV describing the model’s ability to predict developability. **b)** The predictive performance against the independent sequence set describing the models’ ability to generalize beyond the training data. **c)** The generalizable performance was extrapolated to estimate the required number of sequences for the model to perform optimally. Log-log regression was trained with points weighted by the inverse of test loss variance. The error shown represents the propagated error from the standard errors of the parameter estimates.

We also extrapolate how many additional training sequences would be required to achieve performance within the measurement accuracy (experimental variance). For each model, we extrapolated a best-fit line between the log10 test loss and the log10 number of training sequences weighted by the inverse variance for each sample size (Figure 6c). We predict that utilizing the HT assay scores, the number of unique sequences required to obtain optimal performance is 80±40% (P_PK37_, G_SH_, β_SH_) and 81±24% (Sequence and P_PK37_, G_SH_, β_SH_) lower than what would be required when considering sequence information alone, which demonstrates the efficiency of the HT assays to enable developability engineering.

## Discussion

Traditional protein developability measurements are restricted in practical throughput, reducing the number of protein variants that can be reasonably characterized. We evaluated HT assays that genetically encode the POI in a context where the cell’s phenotype is related to the POI’s developability. The on-yeast protease, split GFP, and split β-lactamase assays exhibited their ability to proxy protein developability via prediction of recombinant yield for Gp2 scaffold variants. HT assays increased protein developability knowledge 100-fold (in this study: 400 yield measurements versus predicted yield via 40k HT assay measurements) and enabled analysis of developable sequences. Ligation efficiency for bacterial transformations and the sequencing depth per cost are current capacity limitations. However, future studies utilizing the narrowed set of optimal assay conditions determined in this work could potentially screen millions of unique variants with minimal modifications.

The most useful conditions were determined by comparing the predictive model performance of a traditional developability metric. Only one of six protease assay conditions were utilized in the top model, indicating that other conditions (chemical denaturants, elevated temperature, and alternative protease) were not needed to increase the predictive accuracy of recombinant soluble yield. This may be because the assays were unable to capture alternative stability metrics, or that a single stability metric is sufficient to predict developability. Additional assays may be useful for predicting other traditional developability metrics, such as thermostability. For example, P_TL55_ was found in 5 of 7 top CV models and may aid thermal predictions. The split GFP and split β-lactamase assays were most beneficial when utilizing SH assay scores despite predicting both strain’s yield. We hypothesize SH was able to increase developability resolution over I^q^ in our library by promoting stabilizing disulfide bonds and chaperoning the production of even weakly developable variants.

A nonlinear model was required to convert the split β-lactamase assay HT assay scores to a traditional developability metric. The reference assay evaluated enzymatic activity via minimum inhibitory concentration (MIC) of ampicillin by clonal colony growth on an agar plate^37^. While the exact differences between our measured assay score and the traditional MIC remains unclear, one possible explanation is a decrease in growth rate with increased protein production^46^, lowering the frequency of highly produced variants. Despite the discrepancy, we have shown nonlinear models can extract useful developability information to predict recombinant yield. One assay limitation is the inability to perform direct selection, which is possible for the on-yeast protease and split GFP, based upon the linear model performance. A potential solution to streamline the discovery would be serial direct selections via on-yeast protease and split GFP, followed by a sequenced stratification via the split β-lactamase to increase accuracy.

The Gp2 library (~10^20^) is well beyond the capacity of traditional developability assays that often fail to produce predictive sequence-based models. Utilizing the HT assays we predicted yields 35% closer to experimental accuracy than a one-hot encoded sequence-based model trained on the same sequence set, proving their utility over naïve computational approaches in the vast protein domain. We studied the sitewise amino acid biases based upon predicted yield of 40k unique paratopes, which can be used to design more effective libraries^25,47–50^. However, the analysis utility is limited by multi-site interactions (observed with cysteine) and model accuracy. We believe the increased knowledge will enable more advanced sequence-based models, capable of extrapolating developability to unobserved variants. The efficiency and accuracy of measuring developability proxies via HT assays empowers such models.

We estimate the HT assays will reduce the number of sequences required to produce an optimal predictive model by 80% compared to sequence information alone. Advances in experimental protocol (beyond those evaluated in this study) and alternative model architectures may provide other routes for increased utility. The assays presented in this work have shown the ability to evaluate the developability for a substantially higher number of unique sequences compared to traditional methods. These assays are essentially independent of protein primary function (assuming naïve Gp2 variants tested have no known primary function). Future work will validate the utility of integrating developability assays with discovery and evolution of primary function. Continued improvements of HT assay development may revolutionize the candidate selection process by presorting proteins for ideal developability before the primary function is evaluated, removing a discovery and engineering bottleneck.

## Supporting information

Oligopool.fasta

## Online Methods

See Appended Online Methods PDF.

## Acknowledgements

This work was funded by the National Institutes of Health (R01 EB023339) and a National Science Foundation Graduate Research Fellowship (to A.W.G.). We thank Daniel Woldring for useful feedback on the manuscript. We appreciate support from the University of Minnesota Flow Cytometry Core, University of Minnesota Genomics Center, and the Minnesota Supercomputing Institute (MSI) at the University of Minnesota.

## Author contributions

A.W.G., K.M.M, and B.J.H. designed experiments. A.W.G., K.M.M, S.L., N.N., M.F., and H.P. performed experimental methods. A.W.G., S.M., and B.J.H. analyzed, interpreted, and wrote the manuscript.

## Methods

### Library Generation and Selection

#### Gp2 Insert Preparation

The Gp2 libraries for high-throughput (HT) assays were created via polymerase chain reaction (PCR) overlap extension on oligonucleotides purchased from Integrated DNA Technologies. PCR conditions: 0.02 U/μL Q5 Hot Start High-Fidelity DNA Polymerase (New England Biolabs), 1X Q5 reaction buffer, 200 μM dNTPs, 0.5 μM primers (DNA Table A: 1 & 17), 0.05 μM internal single stranded template (DNA Table A: 2, 3, 4, 11, 15, 16), 0.008 μM degenerate strands for loop 1 (DNA Table A: 5-10), 0.017 μM degenerate strands for loop 2 (DNA Table A: 12-14) diluted to 50 μL with deionized water (dH_2_O). Thermocycling routine: 98 °C for 30 s, (98 °C for 10 s, 59 °C for 30 s, 72 °C for 20 s) x 30 cycles, 72 °C for 120 s. The 300 bp resulting DNA was then gel extracted from a 2% agarose gel and purified via silicon spin column (Epoch Life Sciences) eluting with 30 μL dH_2_O per manufacturer’s instructions.

The constructed library was then amplified for electroporation. PCR conditions: 0.02 U/μL Phusion High-Fidelity DNA Polymerase (New England Biolabs), 1X Phusion HF buffer, 200 μM dNTPs, 0.5 μM primers (DNA Table A: 1 & 17), 15 μL purified template, diluted to 400 μL total volume with dH_2_O. Thermocycling routine: 98 °C for 30 s, (98 °C for 10 s, 66 °C for 30 s, 72 °C for 20 s) x 35 cycles, 72 °C for 120 s. The DNA was concentrated and purified via ethanol precipitation: 40 μL 3 M sodium acetate at pH 5.2 and 1200 μL ethanol was added to the post-PCR product and the mixture was then incubated at 4 °C for 10 min. The insoluble DNA was pelleted via centrifugation at 15,000g for 20 min at 4 °C. The DNA was then washed with 1 mL 70% ethanol in dH_2_O, centrifuged, washed with 1 mL ethanol, centrifuged, aspirated, and dried overnight to R.T. air. The reaction was then resuspended in 30 μL of Buffer E’ (0.5 M sorbitol and 0.5 mM calcium chloride in dH_2_O).

#### Yeast Surface Display Plasmid Preparation

A yeast plasmid display vector pCT from Kruziki et al^1^ was modified to contain a stop codon before the cMyc epitope tag to serve as a negative control (final construct: Aga2-HA-Stop-cMyc, Plasmid Sequence 1). A plasmid expressing the parental Gp2 programmed death-ligand 1 (PD-L1) binding clone E4 was restriction enzyme digested (2 μg plasmid, 20 U BamHI-HF, 20 U PstI-HF, 5U Quick CIP, 1X CutSmart Buffer, diluted to 50 μL with dH_2_O and incubated at 37 °C for 1 hr), extracted from a 2% agarose gel, and purified via silica column eluted with 30 μL of dH_2_O. The motif was inserted via NEBuilder^®^ HiFi DNA Assembly (New England Biolabs): 35 ng of the digested vector, 1.8 ng template DNA (DNA Table B), 5 μL HiFi Master Mix, diluted to 10 μL with dH_2_O was incubated at 50 °C for 20 mins. 2 μL of the reaction mixture was added to 25 μL NEB 5-alpha Competent *E. coli* and transformed according to the manufacturer’s protocol. The transformed cells were plated onto 100 μg/mL ampicillin LB agar plates. A colony was plucked, grown in LB (10 g/L tryptone, 5 g/L yeast extract, 10 g/L sodium chloride) with 100 μg/mL ampicillin, and miniprepped to obtain 50 μg of plasmid. The plasmid was restriction enzyme digested (50 μg plasmid, 200 U BamHI-HF, 200 U PstI-HF, 50 U Quick CIP, 1X CutSmart Buffer, diluted to 500 μL with dH_2_O, incubated at 37 °C for 2 h), and ethanol precipitated. The digested plasmid was reconstituted in 50 μL Buffer E’. The resulting concentration of vector was measured via absorbance with a NanoDrop (Thermo Fisher).

#### Yeast Transformation

The Gp2 gene library was inserted into the yeast display vector (final construct: Aga2–HA–GS linker–Gp2–cMyc) via homologous recombination into *S. cerevisiae* yeast (EBY100) (steps 36-48 in Chao *et al.^2^*) with the following modifications: step 37: inoculated 100 mL of culture, steps 38/41: all (50 μL) of Gp2 insert and 6 μg of digested plasmid was used per 100 mL culture, step 39: included 30% (v/v) PEG 8000 during incubation. Plating a dilution of the transformed cells on selective media estimated the creation of 3.6 x 10^8^ variants of Gp2. Surface display was induced (Step 2^2^) by the introduction of galactose containing media followed by growth at 30 °C overnight.

#### Subsampling Gp2 Library

We chose to subsample the transformed population to increase assay resolution by sampling multiple cells per sequence and performing assays in triplicate. We projected 10 reads per sequence for on-yeast protease and split GFP, and 10 reads per sequence per antibiotic concentration for the split β-lactamase assay, summing to 160 reads per sequence per trial across all ten assays. We found the limiting factors to be the capacity of high-throughput sequencing and bacterial ligation efficiency. Given that an Illumina NovaSeq SP flowcell can achieve 400 x 10^6^ reads per lane for about $3,000, we decided on utilizing two lanes to analyze the 10^6^ sequences to balance information and experimental cost. The realized difference in obtained sequence information is likely due to stochastic sampling leading to a bias in sequence frequencies.

#### Epitope Labeling for Yeast Flow Cytometry

Labeling of the HA and c-Myc epitope tags were used to enrich the frequency of full length Gp2 variants while subsampling from the initial population via fluorescence-activated flow cytometry (FACS). Ten million yeast cells were centrifuged at 5000g for 1 min. The induction media was removed, and the cell pellet was resuspended with 1 mL cold PBSA (1 g/L bovine serum albumin in 1X phosphate-buffered saline (PBS)). The cells were pelleted, aspirated, resuspended in 500 μL PBSA containing 0.5 μg of a chicken-anti-HA antibody (ab9111, Abcam) and 1 μg of a mouse-anti-c-Myc antibody (9E10, Biolegend), and rotated for 30 min at R.T. The cells were then centrifuged, washed with 1 mL cold PBSA, and resuspended in 500 μL PBSA containing 0.5 μg of a goat-anti-chicken AlexaFluor488 antibody (A-11039, Invitrogen) and 1 μg of a goat-anti-mouse AlexaFluor647 antibody (A-21235, Invitrogen) and incubated at 4 °C for 20 min while protected from light. Finally, the cells were centrifuged, washed with 1 mL cold PBSA, and stored as a pellet after centrifugation until sorting. FACS was performed at the University of Minnesota Flow Cytometry Resource facilities. Cells were resuspended at 2 x 10^7^ cells/mL in PBSA and 1 x 10^6^ cells displaying positive 488 (HA) and 647 (c-Myc) signals were collected. Perhaps resulting from multi-vector transformants^3^, 6.7% of the sequences contained a stop codon in the paratope. Additional propagations were performed before completing high-throughput assays to mitigate this issue. Yeast expressing the *GaR* clone, obtained from Kruziki^4^, were added to the subsampled population at an intended ratio of 100 *GaR*: 1 random variant from library (obtained 172:1 via sequencing).

### On-Yeast Protease Assay

#### Reaction Protocol

Dilutions of proteases and yeast were separately prepared on ice. Proteinase K (P8107S, New England Biolabs) was diluted to twice the reaction concentration in PBSA (P_Urea_ was diluted using 3 M urea in PBSA, P_Gdn_ was diluted using 1 M guanidium chloride in PBSA). Thermolysin (V4001, Promega) was reconstituted to 1 mg/mL in 50 mM Tris at pH 8 with 0.5 mM calcium chloride and diluted with PBSA on the day of experiment. Exposure time with protease at reaction temperature was held constant while the concentrations of protease were modified to obtain a roughly equal distribution of FACS gates’ occupancy (Supplemental Figure 1).

Ten million yeast cells expressing the subsampled library were centrifuged at 5000g for 1 min, aspirated, resuspended in 1 mL cold PBSA, centrifuged, resuspended in 50 μL of PBSA, and transferred to a 0.2 mL PCR tube on ice. 50 μL of the diluted enzyme was added to the cells and mixed via pipetting on ice. The enzyme-yeast mixture was placed in a pre-chilled 4 °C PCR block where a preset program heated the mixture to the reaction temperature for 10 min and returned the mixture to 4 °C. Both heating and cooling rates were set to the maximum ramp speed on the Eppendorf Mastercycler Nexus GX2. The enzyme-yeast mixture was then added to 1 mL of cold PBSA and the epitopes were labeled following the protocol used during library subsampling.

The cells were separated via FACS into four populations based upon the c-Myc to HA ratio. The undigested gate (highest c-Myc:HA ratio) was determined by the location of the library in a no-enzyme control. The fully digested gate (lowest c-Myc:HA ratio) was determined by the location of the no-enzyme control where the primary mouse-anti-c-Myc antibody was omitted. The other two gates were drawn to divide the remaining space in half. Collected cells were centrifuged and stored at −80 °C without allowing propagation.

#### Yeast DNA Extraction

Frozen populations were thawed, and the DNA was obtained via Zymoclean Gel DNA Recovery Kit (Zymo Research) following the manufacturer’s protocol. Following the elution into 30 μL of dH_2_O, half of the DNA was mixed with 2 μL ExoI (M0293S, New England Biolabs), 1 μL of Lambda Exonuclease (M0262S, New England Biolabs) and 2 μL of 10 X Lambda Exonuclease Buffer, incubated at 30 °C for 90 min to remove genomic DNA, and 80 °C for 20 min to inactivate the enzymes. The DNA was then purified via silica column purification and eluted with 30 μL dH_2_O.

#### Preparation of DNA for Deep Sequencing

The DNA was prepared for Illumina sequencing and genetically barcoded for population identification by two successive PCR reactions. The first PCR specifically amplified the region of DNA encoding for Gp2: PCR conditions: 0.02 U/μL Q5 High-Fidelity DNA Polymerase (New England Biolabs), 1X Q5 reaction buffer, 200 μM dNTPs, 0.1 μM of 5-forward and 5-reverse primers to add length diversity for sequencing (DNA Table C), 15 μL (half) of the DNA extracted from yeast, diluted to 50 μL total volume with dH_2_O. Thermocycling routine: 98 °C for 30 s, (98 °C for 10 s, 60 °C for 30 s, 72 °C for 20 s) x 16 cycles, 72 °C for 120 s. Unreacted primers were then removed by the addition of 4 U ExoI (37 °C for 30 min, inactivated at 80 °C for 20 min). The second PCR added trial-specific I5 barcode and a gate-specific I7 barcode. PCR conditions: 0.02 U/μL Q5 High-Fidelity DNA Polymerase, 1X Q5 reaction buffer, 200 μM dNTPs, 0.5 μM of forward primer (DNA Table D) and reverse primer (DNA Table E), 1 μL of the DNA from the first PCR, dilluted to 50 μL total volume with dH_2_O. Thermocycling routine: 98 °C for 30 s, (98 °C for 10 s, 67 °C for 30 s, 72 °C for 20 s) x 16 cycles, 72 °C for 120 s. The DNA was purified via agarose gel extraction and quantified via absorbance on a NanoDrop. DNA within the same assay was mixed at the ratio of cells collected during FACS. DNA across assays were evenly mixed for each trial.

### *Split GFP* Assay

#### Creation of GFP_1-10_ Bacterial Production Plasmid

Plasmid pcDNA3.1-GFP(1-10) was a gift from Bo Huang^5^ (Addgene plasmid 70219). The fragment encoding for GFP_1-10_ was isolated via PCR. PCR conditions: 0.02 U/μL Q5 High-Fidelity DNA Polymerase, 1X Q5 reaction buffer, 200 μM dNTPs, 0.5 μM forward and reverse primers (DNA Table F), 1 ng pcDNA3.1-GFP(1-10), diluted to 50 μL with dH_2_O. Thermocycling routine: 98 °C for 30 s, (98 °C for 10 s, 72 °C for 50 s) x 30 cycles, 72 °C for 120 s. The DNA was then purified via silica column.

Plasmid pBAD-His-6-Sumo-TEV-LIC cloning vector (8S) was a gift from Scott Gradia (Addgene plasmid 37507). The plasmid was modified via restriction enzyme digestion (final construct: His_6_-GFP_1-10_, Plasmid Sequence 2). Digestion conditions: 2 μg plasmid, 20 U NheI-HF, 20 U BamHI-HF, 5U Quick CIP, 1X CutSmart Buffer, diluted to 50 μL with dH_2_O and incubated at 37 °C for 1 h. The plasmid was isolated via agarose gel extraction and silica column purification.

GFP_1-10_ was inserted into the pBAD plasmid via NEBuilder^®^ HiFi DNA Assembly (New England Biolabs): 25 ng of the digested vector, 2 ng of GFP_1-10_ encoding DNA, 5 μL HiFi Master Mix, dilluted to 10 μL with dH_2_O and was incubated at 50 °C for 20 min. The assembled plasmid was transformed into NEB 5-alpha Competent *E. coli* as per the manufacturer’s protocol using the ampicillin selection marker.

#### Creation of GFP_11_ Production Plasmid

A pET production plasmid was obtained^6^ and modified to serve as a non-fluorescent control with a stop codon before the C-terminal GFP_11_ (final construct: MAS–Stop–GSGGGGS–GFP_11_, Plasmid Sequence 3).Two rounds of restriction enzyme digestion and HiFi assembly processes were used to complete construction. Digestion 1: 2 μg plasmid, 20 U NheI-HF, 20 U StyI, 5U Quick CIP, 1X CutSmart Buffer, diluted to 50 μL with dH_2_O and incubated at 37 °C for 1 h The plasmid was isolated via agarose gel extraction and silica column purification. HiFi assembly 1: 30 ng plasmid, 2 ng pET-GFP11 gBlock (DNA Table G), 5 μL HiFi Master Mix, diluted to 10 μL with dH_2_O, incubatedat 50 °C for 20 min, and transformed using the kanamycin selection marker. Digestion 2: 2 μg plasmid, 20 U NheI-HF, 20 U BamHI-HF, 5U Quick CIP, 1X CutSmart Buffer, diluted to 50 μL with dH_2_O and incubated at 37 °C for 1 h. The plasmid was isolated via agarose gel extraction and silica column purification. HiFi assembly 1: 30 ng plasmid, 2 ng GFP11-stop insert (DNA Table G), 5 μL HiFi Master Mix, filled to 10 μL with dH_2_O, incubated at 50 °C for 20 min, and transformed using kanamycin selection marker.

#### Ligation of Gp2 Library into GFP_11_ Production Plasmid

The Illumina prepared DNA resulting from the on-yeast protease assay (equal mixture of 6 reaction conditions of trial 1) was used as the source of the Gp2 library for the split GFP assay. The DNA was prepared for ligation via restriction enzyme digest. Digestion conditions: 1.25 μg DNA, 25 U NheI-HF, 25 U BamHI-HF, 1X CutSmart Buffer, filled to 62.5 μL with dH_2_O and incubated at 37 °C for 1 h. The digested DNA was isolated via agarose gel extraction and silica column purification. All pre-ligation gel extractions used Zymoclean Gel DNA Recovery Kits (Zymo Research). The GFP_11_ plasmid was prepared for ligation in a similar process. Digestion conditions: 10 μg DNA, 200 U NheI-HF, 200 U BamHI-HF, 50 U CIP, 1X CutSmart buffer, diluted to 500 μL with dH_2_O and incubated at 37 °C for 1 hour.

Five ligations were required to obtain more than 10^6^ transformed colonies, providing 63% likelihood of sampling any clone of the subsampled library. Ligation conditions: 370 ng vector, 30 ng insert, 10,000 U T4 DNA Ligase (New England Biolabs), 1X T4 Buffer, 1 mM ATP (New England Biolabs), filled to 100 μL with dH_2_O prepared on ice. The reaction was mixed via gentle pipetting and incubated at 22 °C for 15 min, followed by ligase deactivation via incubation at 60 °C for 10 min. The ligated DNA was purified and concentrated into 10 μL dH_2_O via MinElute PCR Purification Kit (Qiagen). The plasmids were transformed into NEB 5-alpha Electrocompetent *E. coli* following the manufacturer’s protocol using 2.5 μL of DNA per 25 μL of cells. An average of 2 x 10^5^ transformed cells was obtained per 100 μL ligation plated on LB agar plates containing 50 mg/L kanamycin. Colonies were scraped from plates and miniprepped to transfer the DNA to production cell lines.

#### Transformation of Split-GFP Production Cells

The GFP_1-10_ plasmid was transformed into T7 Express lysY/I^q^ Competent *E. coli* (I^q^, c3013, New England Biolabs) and SHuffle T7 Express lysY Competent *E. coli* (SH, c3030, New England Biolabs) following the manufacturer’s heat-shock protocol and using the ampicillin selection marker. A single colony from each bacterial strain was plucked and prepared for electroporation: the colony was grown in 100 mL SOB + Amp (2% tryptone, 0.5% yeast extract, 10 mM sodium chloride, 2.5 mM potassium chloride, 10 mM magnesium chloride, 10 mM magnesium sulfate, and 100 mg/L ampicillin in dH_2_O) to an optical density at 600 nm (OD_600_) of 0.5. Unless otherwise stated, strain I^q^ was grown at 37 °C and strain SH was grown at 30 °C. The culture was then placed on wet ice for 15 min and centrifuged (5000g for 10 min). The cells were then resuspended and centrifuged twice with 200 mL of 10% (v/v) glycerol in water. Finally, the cells were resuspended in the residual glycerol before flash freezing with liquid nitrogen and storage at −80 °C.

The Gp2-GFP_11_ plasmids were then electroporated into the prepared competent cells. Frozen cells were thawed on wet ice for 10 min. 20 ng of the plasmid was added to 25 μL of cells and transferred to a cold 1 mm electroporation cuvette. The cells were shocked (2.0 kV, 200 Ω, 25 μF), resuspended in 975 μL SOC (2% tryptone, 0.5% yeast extract, 10 mM NaCl, 2.5 mM KCl,, 10 mM MgCl_2_, 10 mM MgSO_4_, and 20 mM glucose in dH_2_O), and incubated for 1 h. The cells were plated on selective LB agar (containing 100 mg/L ampicillin, and 50 mg/L kanamycin in dH_2_O). Transformations were repeated until >10^7^ colonies were obtained. The colonies were scraped from plates and grown in 100 mL LB with 100 mg/L ampicillin and 100 mg/L kanamycin (LB+Amp+Kan) for two h. Aliquots were created by mixing 1 mL culture with 500 μL glycerol and storing at −80 °C.

#### Split GFP Production Protocol

Frozen aliquots were thawed and grown in 5 mL LB+Amp+Kan overnight. Part of the overnight culture was added to 5 mL fresh LB+Amp+Kan at an OD_600_ of 0.1 and grown for 90 min. Gp2-GFP_11_ production was induced by the addition of 0.5 mM IPTG. For the remainder of split-GFP protocol, both I^q^ and SH strains were grown at 37 °C. Production continued for 2 h, followed by a centrifugation (3000g for 3 min). Cells were then resuspended in 5 mL fresh LB+Amp+Kan and incubated for 1 h to end Gp2-GFP_11_ expression. GFP_1-10_ expression was then induced by adding 2 mg/mL arabinose and production continued for 2 h. Finally, the culture was centrifuged, resuspended in 1 mL cold PBSA, and stored on wet ice.

FACS was used to separate bacterial cells based upon the GFP signal. Background fluorescence was determined by cells containing the stop-GFP_11_ plasmid. The remainder of cells were divided into three equally (log scale) spaced gates. The collected populations were centrifuged (3000g for 10 min) and frozen at −80 °C to inhibit growth. The cells were then thawed and miniprepped to obtain the Gp2-encoding plasmids.

#### Preparation of DNA for Deep Sequencing

DNA was prepared for Illumina as above for the on-yeast protease assay: except for unique primers (DNA Table H) and a 62 °C annealing temperature in the first PCR.

### Split β-lactamase Assay

#### Creation of Production Plasmid

A pET production plasmid was obtained in house and modified via two rounds of restriction digest and HiFi assembly to create a non-functional lactamase control with a stop codon before the second half of the enzyme (final construct: β-lactamase_1-196_-(G_4_S)_2_AS-Stop-GS(G_4_S)_2_-β-lactamase_197-286_, Plasmid Sequence 4). Digestion 1 conditions: 2 μg plasmid, 20 U NdeI-HF, 20 U StyI-HF, 5 U Quick CIP, 1X CutSmart buffer, diluted to 50 μL with dH_2_O and incubated at 37 °C for 1 h), gel extracted, and purified via silica column. HiFi assembly 1: 25 ng plasmid, 2 ng of each insert (Beta-Lac A+B, DNA Table I), 10 μL HiFi Master Mix, filled to 20 μL with dH_2_O, reacted at 50 °C for 15 min, and transformed using kanamycin selection marker. Digestion 2 conditions: 2 μg plasmid, 20 U NheI-HF, 20 U BamHI-HF, 5 U Quick CIP, 1X CutSmart Buffer, filled to 50 μL with dH_2_O and incubated at 37 °C for 1 h), gel extracted, and purified via silica column. HiFi assembly 1: 25 ng plasmid, 2 ng insert (β stop insert, DNA Table I), 10 μL HiFi Master Mix, diluted to 20 μL with dH_2_O, reacted at 50 °C for 15 min, and transformed using kanamycin selection marker. Plasmid for ligation was obtained via miniprep.

#### Split β-lactamase Library Creation and Transformation into Production Cells

The Illumina prepared DNA resulting from the split GFP assay (equal mixture of 2 cell strains from trial 1) was used as the source of the Gp2 library. The DNA was prepared for ligation via restriction enzyme digest. Digestion conditions: 2 μg DNA, 20 U NheI-HF, 20 U BamHI-HF, 1X CutSmart Buffer, diluted to 50 μL with dH_2_O and incubated at 37 °C for 1 h. The digested DNA was isolated via agarose gel extraction and silica column purification. All pre-ligation gel extractions utilized Zymoclean Gel DNA Recovery Kits (Zymo Research). The β-lactamase plasmid was prepared for ligation in a similar process. Digestion conditions: 10 μg DNA, 200 U NheI-HF, 200 U BamHI-HF, 50 U CIP, 1X CutSmart Buffer, diluted to 500 μL with dH_2_O and incubated at 37 °C for 1 h.

Ligations were repeated, using the same conditions as for the split GFP pool above, to obtain more than 10^6^ transformed colonies. Transformed cells were plated on LB agar plates containing 50 mg/L kanamycin. Colonies were scraped from plates and miniprepped to transfer the DNA to production cell lines. The library was transformed into I^q^ and SH following the manufacturer’s heat-shock protocol and using the kanamycin selection marker. Transformations were replicated until more than 10^7^ colonies were obtained. The colonies were scraped from plates and grown in 100 mL LB with 100 mg/L kanamycin for two h. Aliquots were created by mixing 1 mL culture with 500 μL glycerol and storing at −80 °C.

#### Split β-lactamase Assay Protocol

Frozen aliquots were thawed and grown in 5 mL LB+Kan overnight. Part of the overnight culture was added to 5 mL fresh LB+Kan at an OD_600_ of 0.01 and grown for 90 min. The split β-lactamase production was induced by the addition of 0.5 mM IPTG. Production was continued for 2 h at 37 °C (strain I^q^) or 4 h at 30 °C (strain SH). The culture was then divided into 6 x 300 μL wells per concentration of ampicillin in a 96 well plate. 30 μL per well of diluted ampicillin was spiked in to achieve the desired final concentrations. The cultures were then monitored in a Synergy H1 microplate reader (BioTek) with continuous double-orbital shaking and the 600 nm absorbance obtained every five minutes. All wells for a given concentration of ampicillin when the average unnormalized absorbance reached 0.35 were removed from the plate, centrifuged (12,000 g for 3 min), and frozen at −80 °C to stop growth. The cells were then thawed and miniprepped to obtain the Gp2-encoding plasmids.

#### Preparation of DNA for Deep Sequencing

DNA was prepared for Illumina as above for the protease assay except for unique primers (DNA Table J) and a 59 °C annealing temperature in the first PCR.

### High-Throughput Assay Score Calculations

#### Illumina Sequencing and Read Filtering

The prepared DNA from each assay was sequenced via two SP lanes of an NovaSeq 6000 (Illumina) with the help of the University of Minnesota Genomics Center. The first trial for all assays was sequenced in the first lane, and the second and third trials were equally mixed in a second lane after confirming the preliminary success of the first trial.

Sequence analysis was performed using the computational resources of the Minnesota Supercomputing Institute utilizing USearch^7^ to merge, align, filter, denoise, and dereplicate the sequences. Merged reads were clipped to the region between NheI and BamHI prior to quality filtering, where we accepted sequences with less than one expected error based upon the reported quality scores of each nucleotide. A total of 832 x 10^6^ sequences passing filter were obtained (Trial 1: 434 x 10^6^, Trial 2: 211 x 10^6^, Trial 3: 188 x 10^6^).

A contamination of DNA encoding for Gp2 with a PD-L1 binding paratope and framework mutations was experimentally confirmed in the on-yeast protease assay sequencing primer stocks. Thus, these sequences were removed from the on-yeast assays but were included in the split GFP and split β-lactamase assays as the sequences were obtained from the on-yeast DNA.

Beyond the physical contamination, an average of 56 x 10^6^ unique sequences was obtained per trial, well beyond the expected max diversity of 1 x 10^6^. We hypothesize the “true” sequences to have high observation frequency (most number of reads) and contain highly different sequences compared to “false” sequences (due to subsampling of theoretical library, it is unlikely to see two very similar sequences). We denoised each trial independently utilizing the UNOISE^8^ algorithm with observation minimums chosen for computational efficiency (Trial 1: 100 total reads, Trial 2 and 3: 50 total reads). A total of 294,644 unique sequences observed in all three trials were obtained from denoising. We then mapped the filtered false sequences to the true sequence via 97% genetic similarity. Finally, 204,173 unique sequences for *CC+, CC−*, and *Stop* were isolated via requiring 100% genetic match of the conserved portion of Gp2. *CC+* was identified via two cysteines located at each end of loop 1 (positions 7 & 12), whereas other sequences were classified *CC−. Stop* sequences contained at least one stop codon located inside either loop, but otherwise matched library design. No sequences containing synonymous codons were observed during the measurement of recombinant yield.

#### On-yeast protease and Split GFP Assay Score Calculation

The four collection gates in the FACS based assays were drawn to bin cells via hypothesized developability. Thus, we defined an assay score which correlates to the relative position of a sequence. To increase resolution, we collected an average of 6.7X (on-yeast protease) and 7.9X (split-GFP) the hypothesized diversity of cells per trial and assigned a score correlating to the average cell location.

For each population, the read frequency of every sequence was converted to the number of cells collected via FACS (Equation M1).

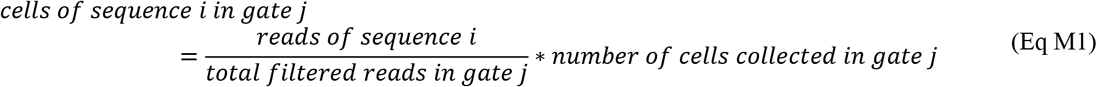

The assay score for a sequence was calculated by assigning each gate a score [0, 1/3, 2/3, 1] and determining the cell-averaged score (Equation M2). For on-yeast protease, 1 was given to full length sequences and 0 was given to fully digested sequences. For split GFP, 0 was given to no detected GFP signal and 1 was given to the highest amount of GFP signal.

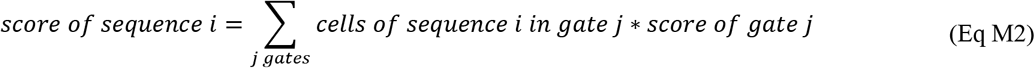

The final assay score was determined by the average score for a sequence in each trial. Sequences without reads in at least one gate per trial were removed from the dataset.

#### Split β-lactamase Assay Score Calculation

We aimed to assign an assay score that would correlate to the total activity of *β*-lactamase enzyme in each cell. We assumed that cells with active enzyme grown in ampicillin will retain the ability to grow and divide (and thus increase DNA frequency), whereas cells with inactive enzyme grown in ampicillin will stop growth (and thus prevent any increase in DNA frequency). To increase resolution, we chose ampicillin concentrations that produced approximately 10%, 30%, and 60% of uninhibited growth for each cell strain. Briefly, we estimated the max growth rate and determined the extra number of doublings required to reach a given concentration. Assuming all cells are growing with no ampicillin, the relative number of dividing cells can be determined by the initial number of cells. The assay score for each sequence was determined by the relative change in read frequency with increasing ampicillin concentrations. For simplicity, the ampicillin concentrations were assigned to [0, 1, 2, 3] where 0 represented the no-ampicillin control and 3 represented the highest ampicillin concentration.

The final assay score was determined by the average score for a sequence in each trial. Sequences without a read in the no-ampicillin population in each trial were removed from the dataset. To scale the assay scores within the range [0,1], scores for *CC+* and *CC−* sequences (not including the independent Test sequences to prevent data leaking) were normalized via scikit-learn’s quantile transformer with a normal output distribution followed by a minmax scaler.

### Dot Blots to Quantify Expression

#### Creation of Production Plasmid

A pET production plasmid was obtained in-house and modified via restriction digest and HiFi assembly to create a His_6_-less negative control with a stop codon placed between the restriction sites. (final construct: V5-AS-Stop-GS-His_6_, Plasmid Sequence 5). Digestion conditions: 2 μg plasmid, 20 U NheI-HF, 20 U BamHI-HF, 5 U Quick CIP, 1X CutSmart Buffer, diluted to 50 μL with dH_2_O and incubated at 37 °C for 1 h), gel extracted, and purified via silica column. HiFi assembly 1: 25 ng plasmid, 2 ng insert (DNA Table K), 10 μL HiFi Master Mix, diluted to 20 μL with dH_2_O, incubated at 50 °C for 15 min, and transformed utilizing kanamycin selection marker. Plasmid for ligation was obtained via miniprep.

#### Dot Blot Library Creation and Transformation into Production Cells

DNA encoding for Gp2 variants for the dot blots were obtained from two sources: i) the Illumina prepared DNA resulting from the split β-lactamase assay (equal mixture of the no-ampicillin population from both cell strains in trial 1). ii) an oligopool (Oligopool.fasta, Twist Bioscience) encoding for the 2,000 most frequently observed Gp2 variants found in all 10 assays of trial 1. The oligopool was amplified to create double stranded DNA: PCR conditions: 0.02 U/μL Q5 Hot Start High-Fidelity DNA Polymerase (New England Biolabs), 1X Q5 reaction buffer, 200 μM dNTPs, 0.5 μM primers (DNA Table L), 10 ng of the oligopool resuspended at 1 ng/μL in EB (Epoch Life Science), diluted to 50 μL total volume with dH_2_O. Thermocycling routine: 98 °C for 30 s, (98 °C for 10 s, 61 °C for 30 s, 72 °C for 20 s) x 12 cycles, 72 °C for 120 s. The amplified oligopool was silica column purified eluting with 50 μL dH_2_O.

The DNA from each source was separately prepared for ligation via restriction enzyme digest. Digestion conditions: 2 μg DNA, 20 U NheI-HF, 20 U BamHI-HF, 1X CutSmart Buffer, filled to 50 μL with dH_2_O and incubated at 37 °C for 1 h. The digested DNA was isolated via agarose gel extraction and silica column purification. All pre-ligation gel extractions used Zymoclean Gel DNA Recovery Kits (Zymo Research). The His_6_ production plasmid was prepared for ligation in a similar process. Digestion conditions: 2 μg DNA, 20 U NheI-HF, 20 U BamHI-HF, 1X CutSmart Buffer, diluted to 50 μL with dH_2_O and incubated at 37 °C for 1 hr.

Ligations were repeated to obtain more than 10^3^ transformed colonies, to obtain a feasible testable diversity of sequences. Ligation conditions: 37 ng vector, 3 ng insert, 1,000 U T4 DNA Ligase (New England Biolabs), 1X T4 Buffer, 1 mM ATP (New England Biolabs), filled to 10 μL with dH_2_O prepared on ice. The reaction was mixed via gentle pipetting and incubated at 22 °C for 15 min, followed by ligase deactivation via incubation at 60 ° C for 10 min. The plasmids were heat-transformed into NEB 5-alpha Competent *E. coli* following the manufacturer’s protocol and using 2.5 μL of DNA per 25 μL of cells. Transformed cells were plated on LB agar plates containing 50 mg/L kanamycin. Colonies were scraped from plates and miniprepped to transfer the DNA to production cell lines. The library was transformed into I^q^ and SH following the manufacturer’s heat-shock protocol and using the kanamycin selection marker. Transformations were replicated until more than 10^3^ colonies were obtained. Single colonies were grown in 1 mL LB+Kan for two hours in separate wells of a deep 96 well plate. Aliquots were created by mixing 1 mL culture with 500 μL glycerol and storing at −80 °C.

#### Identifying Plate Location of Gp2 Variants

We appended genetic barcodes representing the plate, row, and column via PCR while simultaneously preparing the sequences for Illumina sequencing. PCR 1 conditions: 0.02 U/μL Q5 High-Fidelity DNA Polymerase (New England Biolabs), 1X Q5 reaction buffer, 200 μM dNTPs, 0.5 μM of a row-specific forward primer (DNA Table M FApETV5N50X), 0.5 μM of a column-specific forward primer (DNA Table M RApETN7XX), 1 μL bacterial culture, filled to 20 μL total volume with dH_2_O. Thermocycling routine: 98 °C for 5 min, (98 °C for 10 s, 61 °C for 30 s, 72 °C for 20 s) x 16 cycles, 72 °C for 120 s. The DNA for each plate was pooled at equal volume (2 μL) and the unreacted primers were then removed by the addition of 8 U ExoI (37 °C for 30 minutes, inactivated 80 °C for 20 min). The second PCR added a plate specific barcode. PCR conditions: 0.02 U/μL Q5 High-Fidelity DNA Polymerase, 1X Q5 reaction buffer, 200 μM dNTPs, 0.5 μM of forward primer (DNA Table D) and reverse primer (DNA Table E), 1 μL of the DNA from the first PCR, filled to 50 μL total volume with dH_2_O. Thermocycling routine: 98 °C for 30s, (98 °C for 10 s, 67 °C for 30 s, 72 °C for 20 s) x 16 cycles, 72 °C for 120 s. The DNA was isolated via agarose gel extraction and quantified via NanoDrop. DNA across plates and cell strain were equally mixed and sequenced via Illumina iSeq, aiming to obtain ~1,000 reads per well.

Sequence analysis was performed using the computational resources of the Minnesota Supercomputing Institute utilizing USearch^7^ to merge, align, filter, denoise, and dereplicate the sequences. Merged reads were clipped to the region between NheI and BamHI prior to quality filtering, where we accepted sequences with less than one expected error based upon the reported quality scores of each nucleotide. To identify single-variant wells, the most abundant sequence had to have >100 reads, the next most sequence had to have <100 reads, and the top sequence had to occupy > 80% (DNA from β-lactamase) or > 40% (DNA from oligopool, changed based upon resequencing previously identified variants) of the total reads for a well. Sequences obtained from dot blot sequencing were genetically paired (requiring 100% matching) with sequences from HT assays.

#### Preparation of Protein Standard

*GaR* was separately ligated into the production vector (final construct V5-*GaR*-His_6_) and transformed following the same protocol used for the library of Gp2 variants, with the difference of transforming the post-ligation product directly into I^q^ via heat-shock transformation. A scrape from the frozen stock of *GaR* cells was grown in 5 mL LB+Kan overnight. Part of the overnight culture was added to 200 mL fresh LB+Kan at an OD_600_ of 0.1 and grown for 90 min. The protein production was induced by the addition of 0.5 mM IPTG. Production was continued for 2 h at 37 °C followed by centrifugation (3,000 g for 15 min) and freezing of the cell pellet at −80 °C overnight. The pellet was thawed by the addition of 2 mL lysis buffer: 1 mg/mL lysozyme (L6876, Millipore Sigma), 10 U/mL benzonase nuclease (E1014, Millipore Sigma), protease inhibitor pellet (A32953, Thermo Fisher Scientific), 20 mM sodium chloride, 2 mM magnesium chloride, 25 mM imidazole, 5 mM 3-[(3-cholamidopropyl)dimethylammonio]-1-propanesulfonate hydrate (CHAPS), 5% (v/v) glycerol, 50 mM (4-(2-hydroxyethyl)-1-piperazineethanesulfonic acid) (HEPES) in dH_2_O at pH 8.0. The lysate and lysis buffer were shaken at 37 °C for 1 h to promote enzymatic activity. The soluble lysate was isolated via centrifugation (15,000g for 10 min) and filtered through a 0.22 μm membrane. *GaR* was isolated via immobilized metal affinity chromatography utilizing HisPur Cobalt Resin (89964, ThermoFisher Scientific); wash buffer: 500 mM sodium chloride, 20 mM HEPES, 20 mM imidazole, pH 7.4; elution buffer: as in wash but 500 mM imidazole. The protein was desalted on a PD-10 column (Fisher Scientific, eluted into 0.5 M sodium chloride, 20 mM HEPES, pH 7.4). The identity and purity were confirmed via matrix-assisted laser desorption/ionization (MALDI) and polyacrylamide gel electrophoresis (PAGE), and concentration was determined via 280 nm absorbance on a NanoDrop. The protein was diluted into 4 standard concentrations (103 ng/μL, 52 ng/μL, 26 ng/μL, and 13 ng/μL) and flash frozen in aliquots via liquid nitrogen and stored at −80 °C. On the day of use, the aliquot was thawed and 25 μL of protein was mixed with 25 μL of denaturing buffer (1 g/L SDS, 500 mM sodium chloride, 20 mM imidazole, 20 mM HEPES, pH 7.4) and incubated at 70 °C for 5 min.

#### Production of Gp2 Library for Dot Blot

Frozen cells from deep well 96-well plates were scraped and seeded into 500 μL/well fresh LB+Kan and grown overnight (I^q^ was grown at 37 °C and SH was grown at 30 °C for the entire production). The following day, 25 μL/well of overnight culture was added to 1 mL/well of fresh LB+Kan and grown for 90 min. The protein production was induced by the addition of 0.5 mM IPTG (diluted in LB+Kan to add 100 μL/well). Production was continued for 2 h (I^q^) or 4 h (SH) followed by centrifugation (3,000g for 5 minutes) and freezing of the cell pellet at −80 °C overnight. The pellet was thawed by the addition of 100 μL/well lysis buffer (only change is 0.1 mg/mL lysozyme) and shaken at 37 °C for 1 hour. The plates were centrifuged (3,000g for 5 min) and 25 μL/well of the soluble fraction was added to 25 μL/well of denaturing buffer Protein lysates from SH were diluted an additional 5X in denaturing buffer to ensure signals were within the range of standards. The plates were incubated at 70 °C for 5 min to ensure denaturation and full accessibility of the His_6_ tag.

#### Dot-Blot Protocol

A section of 0.2 μm pore polyvinylidene fluoride (PVDF, 1620177, BioRad) was cut to size and placed in a box (15.2 cm × 10.2 cm × 3.2 cm, Z742094, Sigma Aldrich). The membrane was soaked in 50 mL methanol for 30 s, followed by 50 mL dH_2_O for 2 min. Finally, the membrane was equilibrated in 50 mL TBST (0.05% v/v Tween 20 in tris-buffered saline (TBS)) for 5 min. The membrane was then placed on a TBST soaked filter paper and padded dry with a Kimwipes™. Using a multichannel pipet, 2 μL/well of protein samples were added to the membrane and allowed to fully absorb. The membrane was then transferred to a dry filter paper and placed in a fume hood for 30 min until dry. The membrane was then placed back in the box with 50 mL blocking solution (5% (w/v) nonfat dry milk in TBST) and rocked overnight at 4 °C. The membrane was then labeled with 50 mL of 0.2 μg/mL anti His_6_-HRP (ab1187, Abcam) in blocking solution for 30 minutes at room temperature. Excess antibody was washed via 3 washes of 50 mL TBST for 10 min at room temperature. The membrane was then soaked in 25 mL of SuperSignal™ West Pico PLUS Chemiluminescent Substrate (ThermoFisher) for 5 min. Then membrane was then placed inside a transparency and exposed 10-30 s on a ChemiDoc MP Imaging System (BioRad).

#### Quantification of Chemiluminescent Intensities

Intensity measurements were quantified utilizing Fiji^9,10^. The average intensity of a constant diameter of a circular region of interest for each lysate impression was recorded. Ten randomly chosen background locations were also measured and subtracted from the intensity measurements. A row of standards (4 concentrations, each concentration in triplicate) was used to generate a linear standard curve from average intensity to yield (mg/L). To correct for non-specific binding of *E. coli* lysate proteins (not present in standard curve), the 75^th^ percentile value of yield for wells containing *Stop* sequences in each trial was subtracted. Sequences with negative corrected yields were set to 0 mg/L. The final yield for model evaluation was reported as the average of three yield measurements, grown from separate starter cultures on different days. *GaR* was tested on each plate for both cell types to obtain an estimate of variance. It was observed that day-to-day coefficient of variation (I^q^: 43%, SH: 77%) was higher than plate-to-plate variation (I^q^: 20%, SH: 25%).

### Identification of HT Assay Predictiveness

#### Code Availability

Python scripts used for deep-sequencing and model evaluation, as well as datasets to train, evaluate, and plot predictive performance are available at https://github.com/HackelLab-UMN/DevRep.

#### Sequence Encoding

To create models with the amino acid sequence, we only considered the amino acids in the modified paratope loops. To conserve possible interactions with the first/last position and the conserved residues of the protein, gap characters were placed in the middle of loops during sequence alignment. The gap character was treated as a 21^st^ amino acid during the one-hot encoding of the sequence.

#### Cross-Validation Performance

A set of 195 unique Gp2 variants contained measured HT assay scores in all 10 assay conditions, and a yield in at least one of the strains. We performed 10 x 10 repeated K-fold cross-validation to determine which of the 1,023 combinations of HT assay conditions predicted the “left-out” set of sequences’ yield with the least error. Each HT assay combination was evaluated for predictive performance on four different model architectures summarized in Table M1. We utilized the Hyperopt^11^ library to determine the optimal hyperparameters for each architecture. We allowed 50 trials (or a maximum of 24 hours of computational time for FNN) and recorded the trial with the lowest predictive error.

**Table M1:**
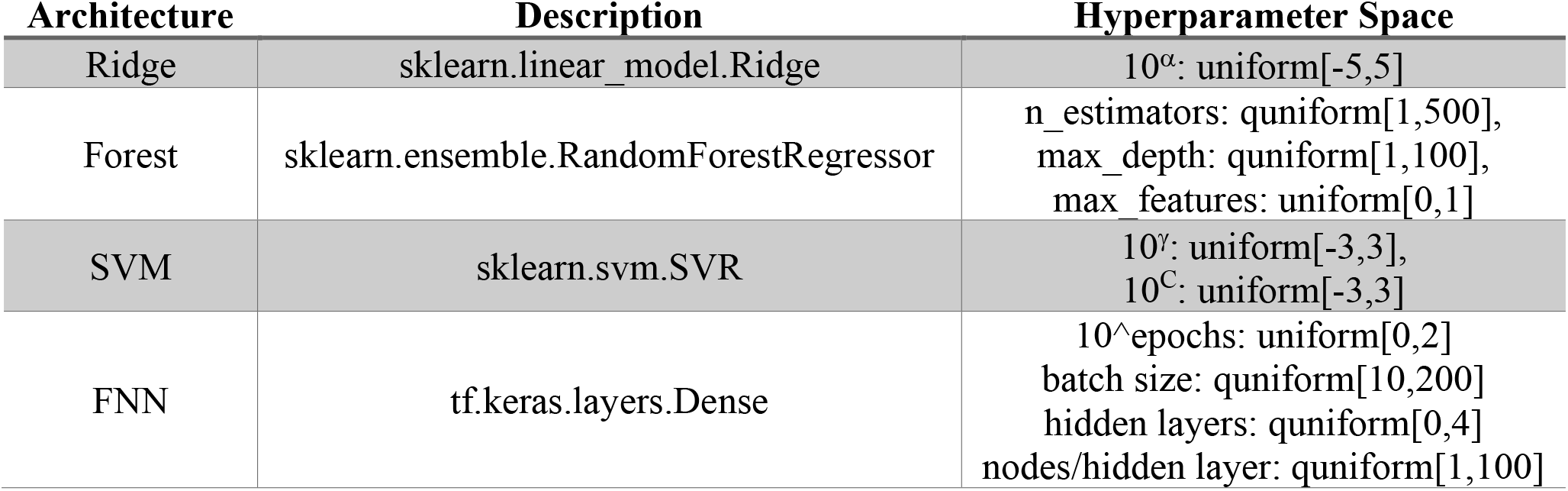
Description of model architectures utilized when evaluating HT assay predictive performance. “uniform” and “quniform” refers to stochastic search spaces defined in the Python *hyperopt* library^12^.

#### Test Performance

When evaluating performance on the independent test sequences, the best model architecture and hyperparameters were chosen by cross-validation, but the weights for the model were refit utilizing the entire cross-validation training set. The independent test set was not used in training data transformations or models.

#### Correlation Feature Selection (CFS)

CFS identifies the optimal feature set by maximizing the relationship between features (x, HT assays) and target (y, yield) while minimizing the inter-feature relationships^13^. We calculated the CFS for every set (Sx) of 1023 HT assay combinations. We defined the relationship (r) as the absolute value of Spearman’s rank correlation coefficient (ρ) or the mutual information (MI) to capture linear and nonlinear relationships (Equation M3).

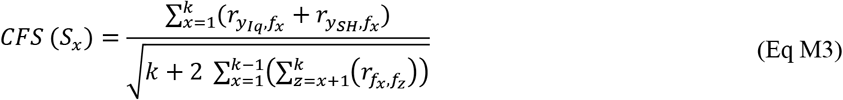

#### Subsampling Training Data

When evaluating the predictive performance of assays with varying number of training datapoints, we bootstrapped the dataset for cross-validation ten times. Each random dataset had separately optimized architectures and hyperparameters determined by cross-validation. Due to the computational constraints, FNN architecture was not evaluated when subsampling the training dataset.

#### Propagation of Uncertainty

Calculations involving propagation of uncertainty for predicted sample size were performed using the *uncertainties*^14^ Python package.

---

**Supplementary Figure 1.**
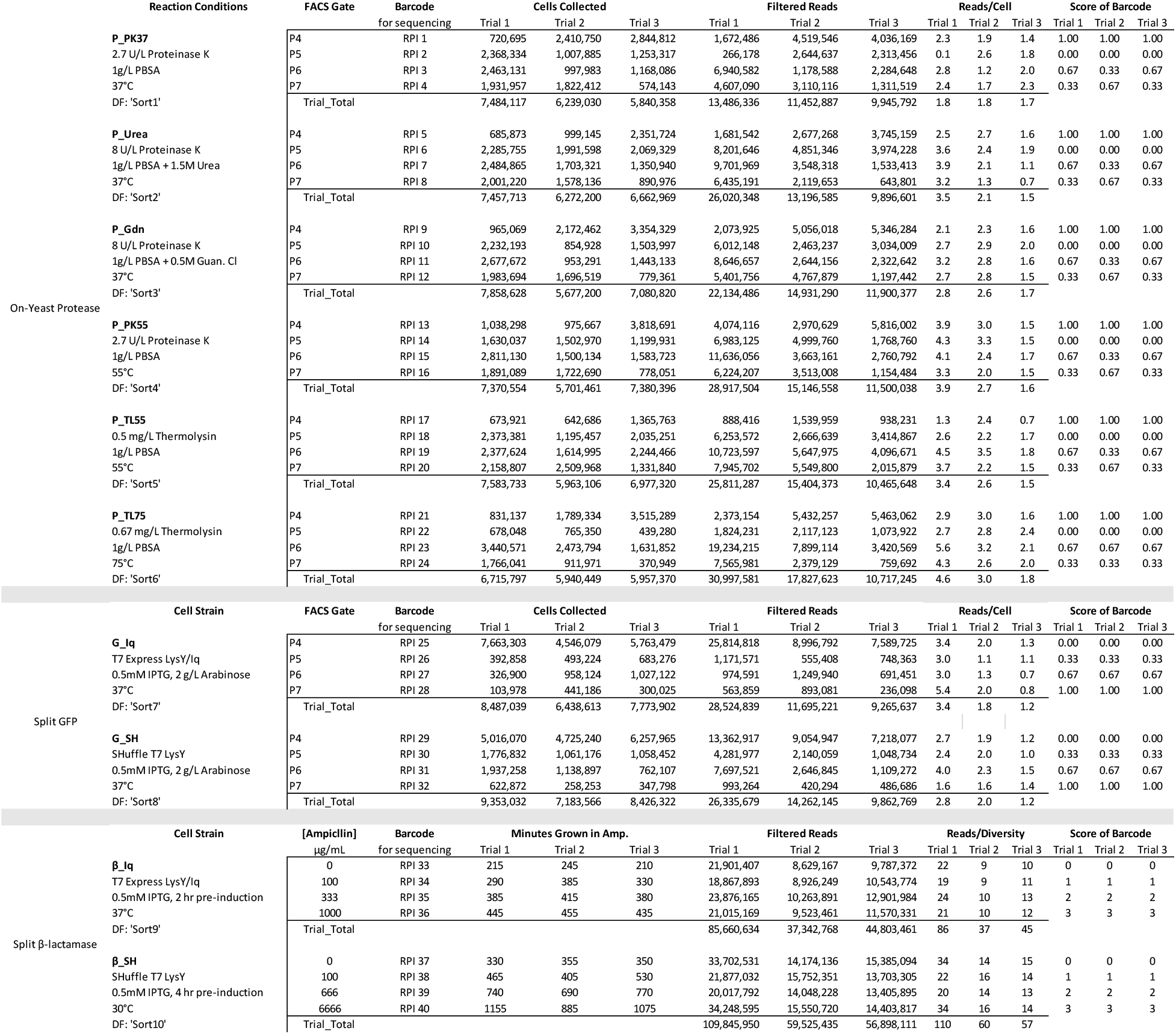
High-throughput (HT) Developability Assay Experimental Results. For each HT assay class, the varying conditions are listed along with the short-hand code and the name of the relevant DataFrame (DF) columns which store assay scores. On-yeast protease and split GFP FACS gates represent polygons named during experimentation. The split β-lactamase assay included a pre-induction of the enzyme-POI fusion before introduction of ampicillin. The *Minutes Grown in Amp*. represents the amount of time until the culture begins to exit logarithmic growth phase based upon preliminary assay testing. *Filtered Reads* represents the number of merged and aligned paired-end reads matching design and was used as the denominator when calculating read frequency. The scores for each barcode were utilized when calculating the cell-averaged score (on-yeast protease and split GFP) or when calculating the change in frequency as a function of ampicillin concentration (slope uses scores not actual concentration).

**Supplementary Figure 2.**
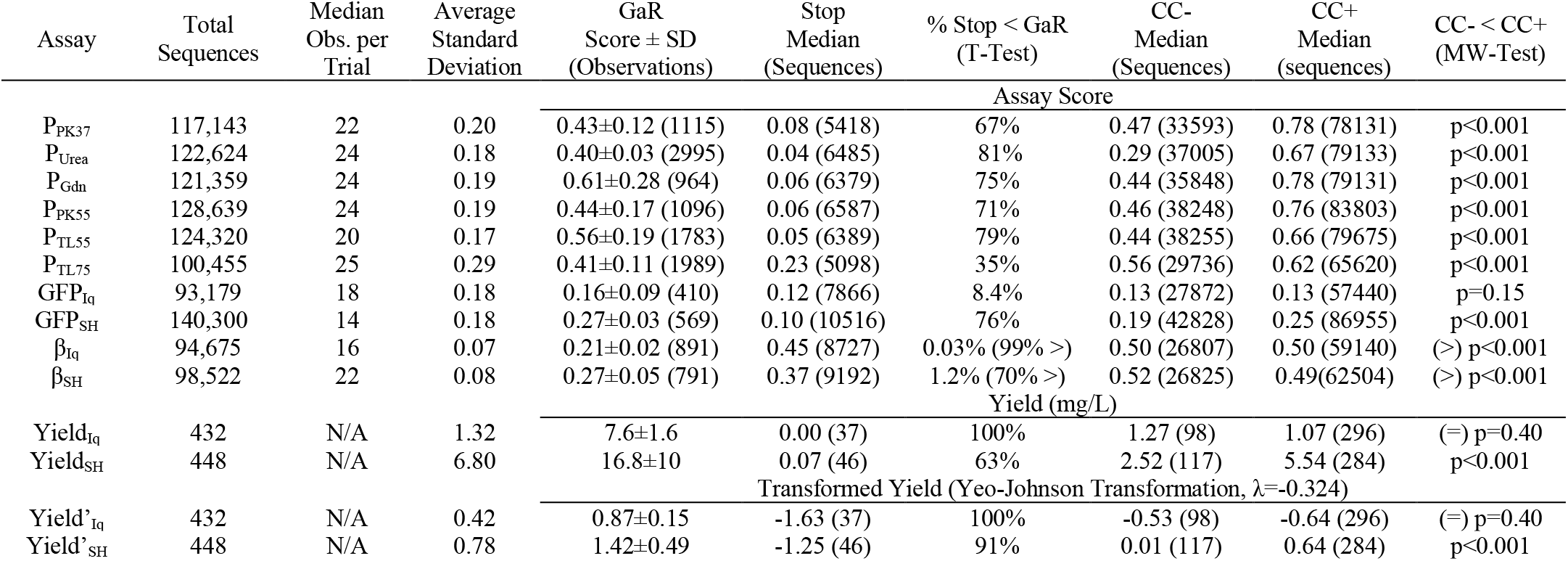
Tabulated Gp2 Library Developability Performance. *Total Sequences*: all Stop, CC−, and CC+ sequences observed in all 3 trials per assay. *Median Observations per Trial* for on-yeast Protease and Split GFP assays measured observations as the number of cells collected in all gates per trial. Split β-Lactamase measured observations as the number of reads in the no-ampicillin control due to anticipated loss of sequences which fail to replicate in antibiotic-containing conditions. *Average Standard Deviation* represents the square root of the mean (by unique sequence) of experimental variance across 3 independent trials. *% Stop < GaR* is the percentage of unique stop codon sequences for which the assay score was significantly lower than the GaR score (one-way student’s t-test, p<0.05). For β_Iq_ and β_SH_, % Stop > GaR is also displayed. *CC− < CC+* tests the hypothesis that sequences with cysteines at positions 7 and 12 significantly increase assay score (one-way Mann-Whitney U test). For β_Iq_ and β_SH_, the opposite hypothesis, CC− > CC+, is displayed. For YieldIq, the two-sided Mann-Whitney tested is displayed.

**Supplementary Figure 3.**
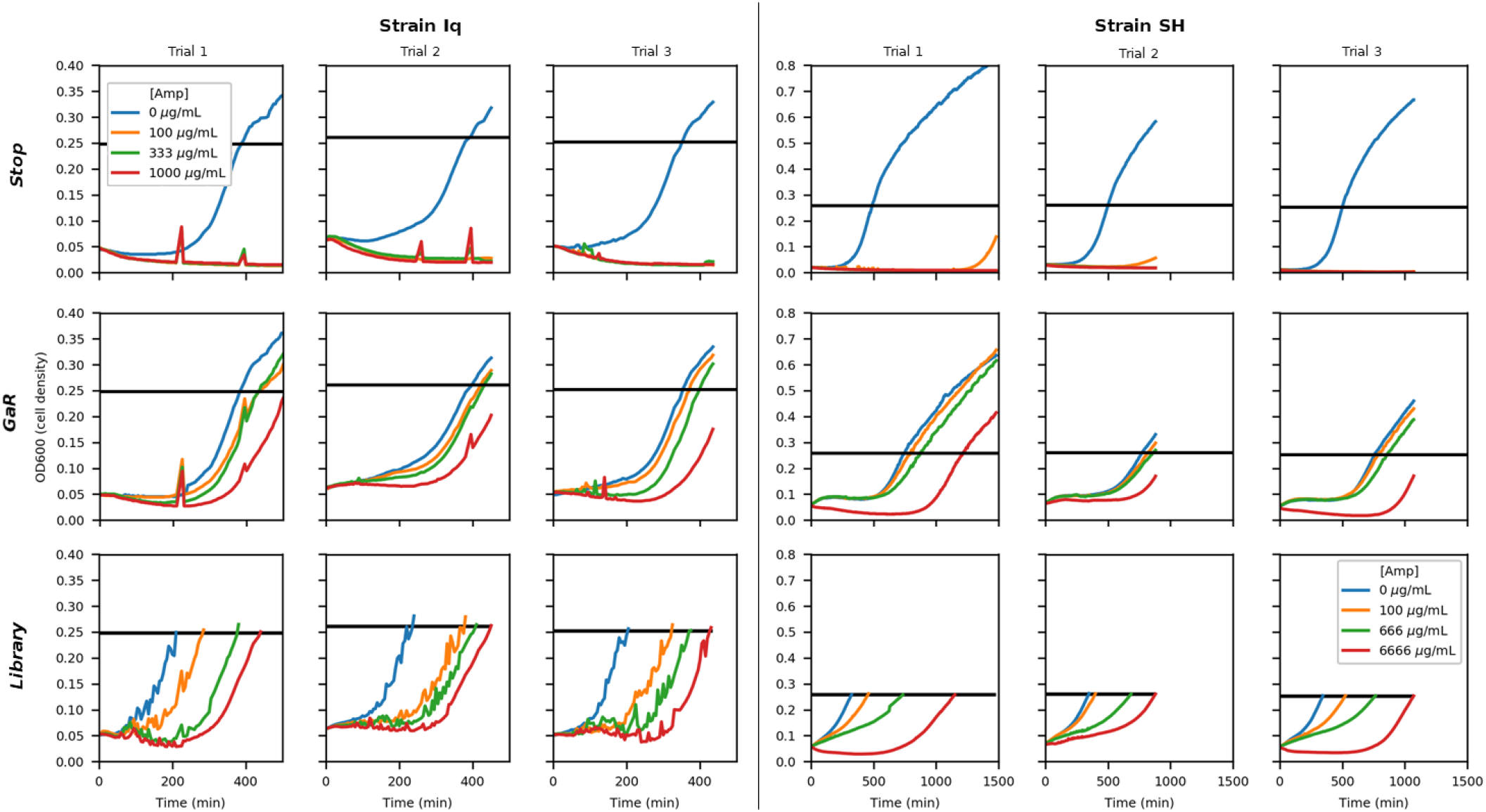
Split β-lactamase Growth Curves of Unmixed *Stop* and *GaR* and mixed Library. The growth curves of the control sequences grown in individual wells and the pooled library of cells. The black bar represents the cell density at which the library was collected for sequencing.

**Supplementary Figure 4.**
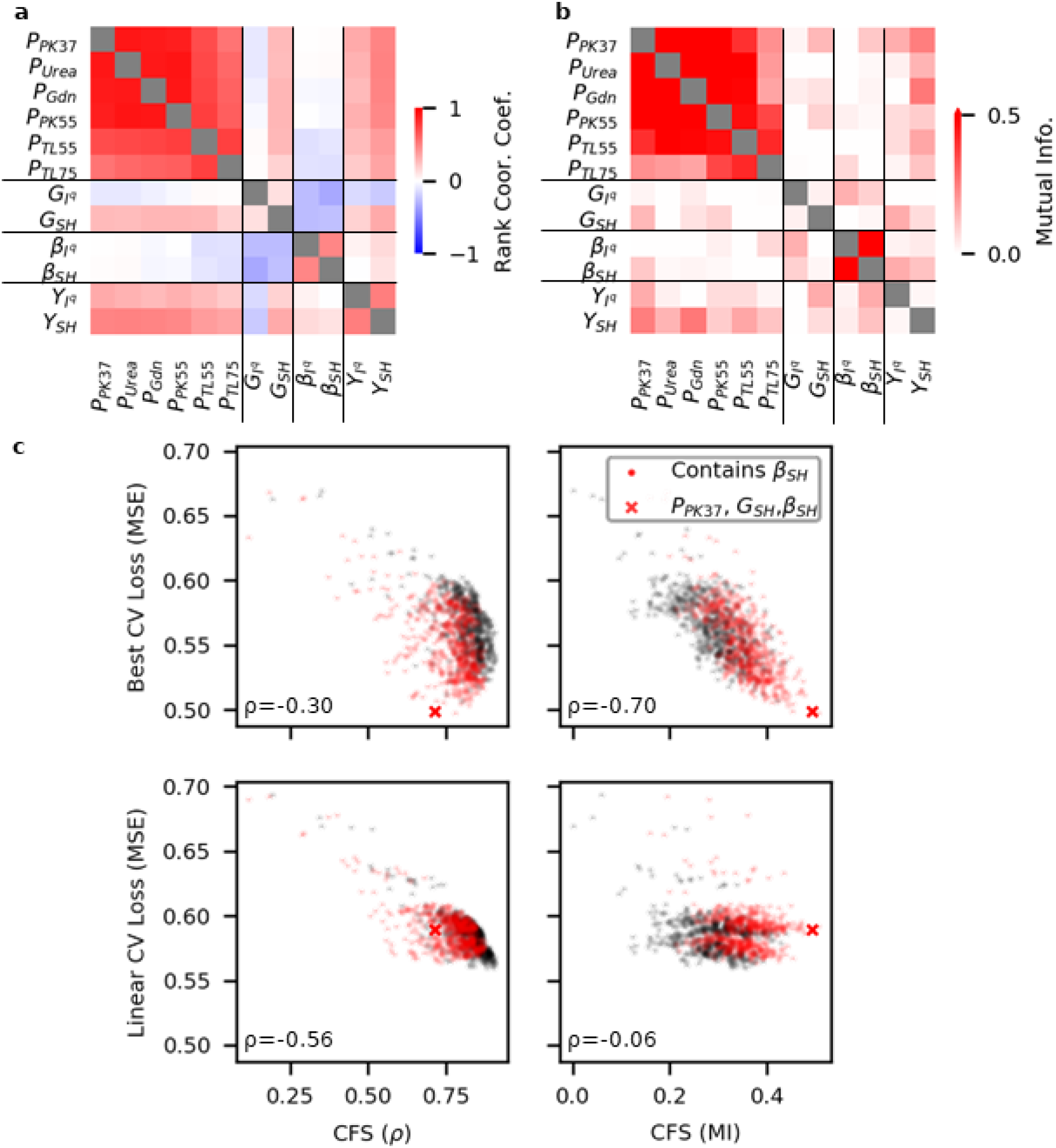
Correlation feature selection (CFS) confirms the selection of most predict HT assay conditions. **a)** The Spearman’s rank correlation coefficient (ρ) and **b)** the mutual information (MI) between HT assays and yield. **c)** Scatter plot of CFS as calculated by p (left) or MI (right) versus predictive loss for the best model architecture (top, best of: Ridge, SVM, Forest, FNN) or a linear model (bottom, Ridge) for the 1023 combinations of HT assays. The ρ between CFS and model loss is presented in the lower-left corner.

**Supplementary Figure 5.**
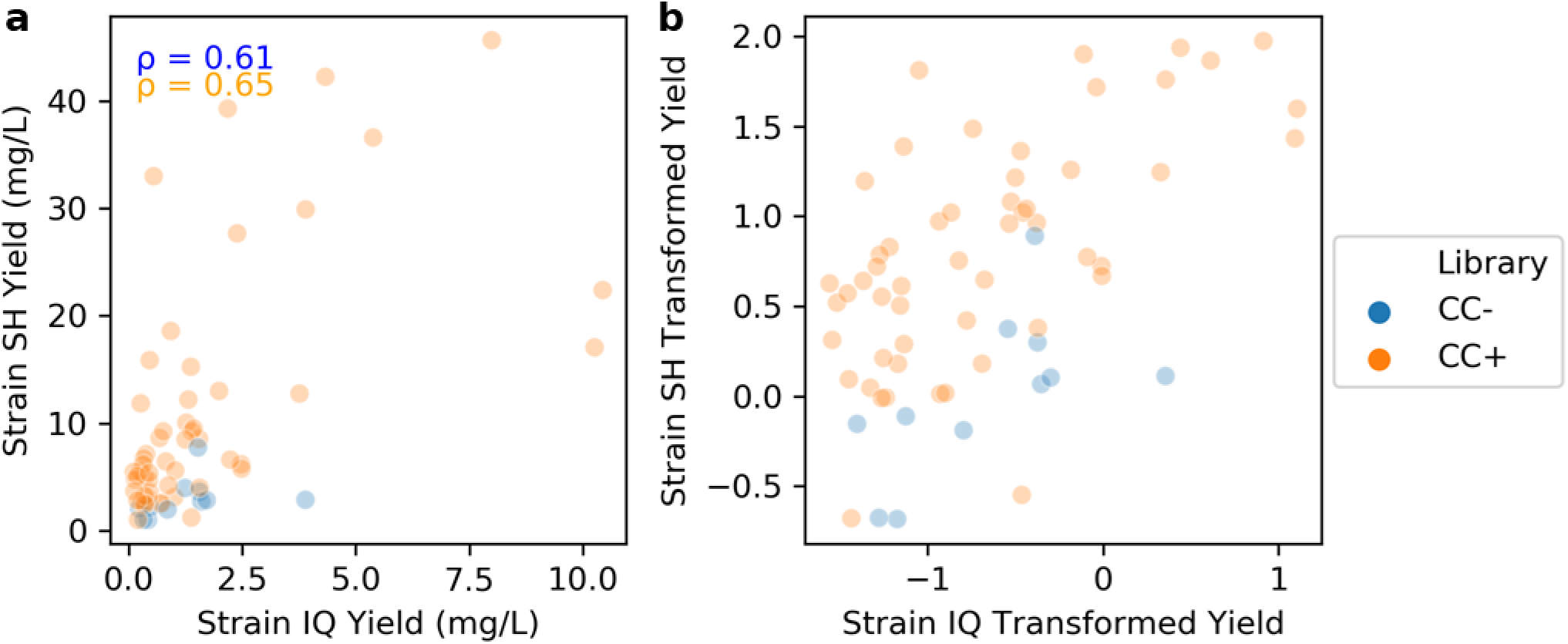
Correlation of yields between cellular strains suggests shared information. **a)** Scatter plot of recombinant soluble yield of 64 (*CC+ orange*: 53, *CC−, blue*: 11) unique Gp2 mutants produced in both bacterial strains as measured by chemiluminescent dot blot. The Spearman’s rank correlation coefficient (ρ) is shown for each sequence class in the upper left corner. **b)** Scatter plot of the same sequences but utilizing the transformed yield used during HT assay predictions (see Supplementary Figure 6).

**Supplementary Figure 6.**
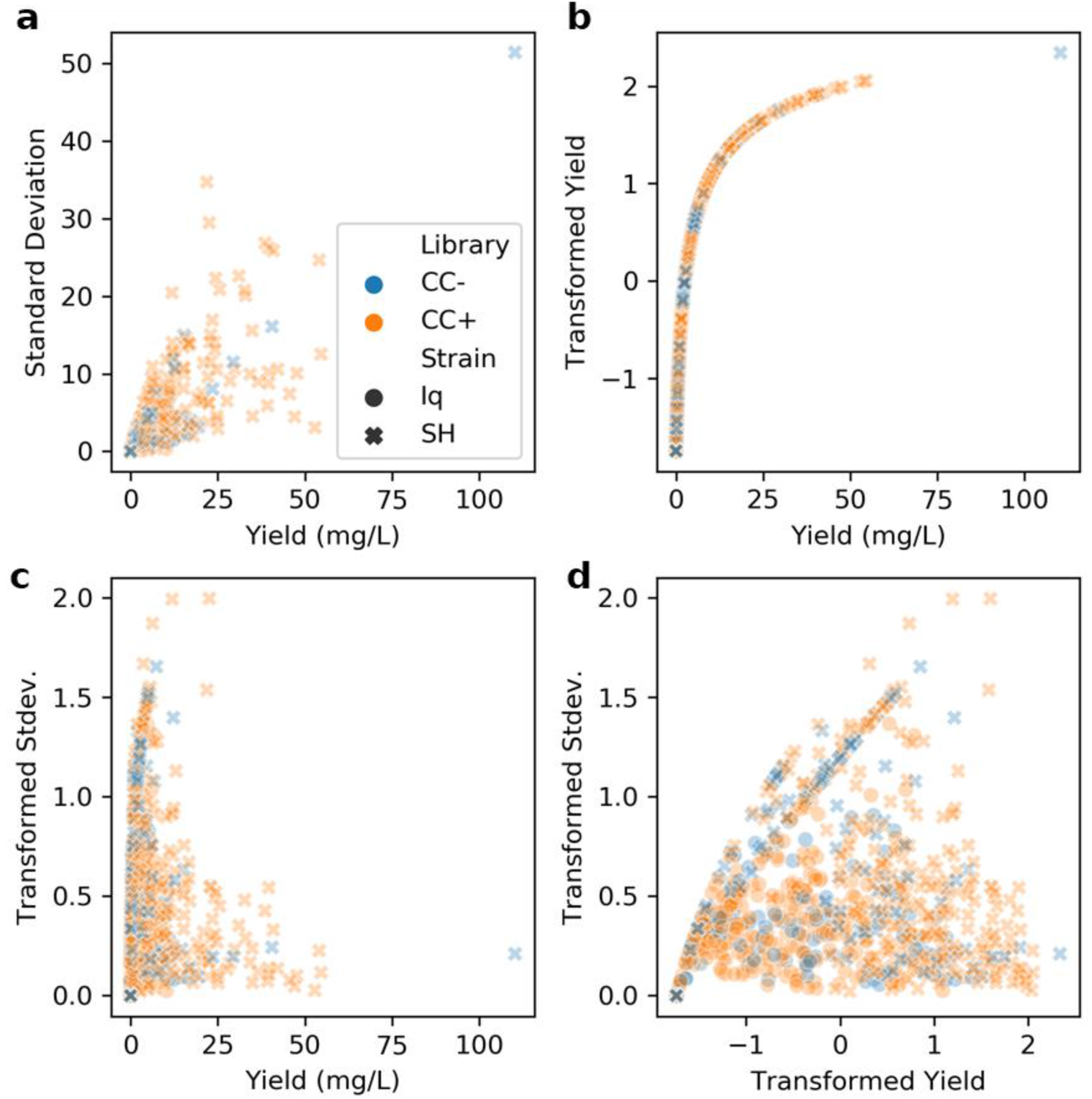
Yeo-Johnson power transformation and normalization of yield. Transformation was performed to remove heteroscedasticity (increasing experimental error with increasing yield). A single transformation parameter (λ = −0.324) was trained using only those sequences utilized in cross-validation for both libraries (*CC+*: orange, *CC−*: blue) and production strains (Iq: circle, SH: cross). **a)** Scatter plot of the yield for unique Gp2 sequences versus the trial-to-trial (n=3) standard deviation showing an increasing trend. **b)** Scatter plot of the measured yield via calibration curve of the chemiluminescent dot blot versus the transformed yield used for model evaluation. **c,d**) Scatter plots showing the removal of increasing experimental error with increasing transformed yield.

**Supplementary Figure 7.**
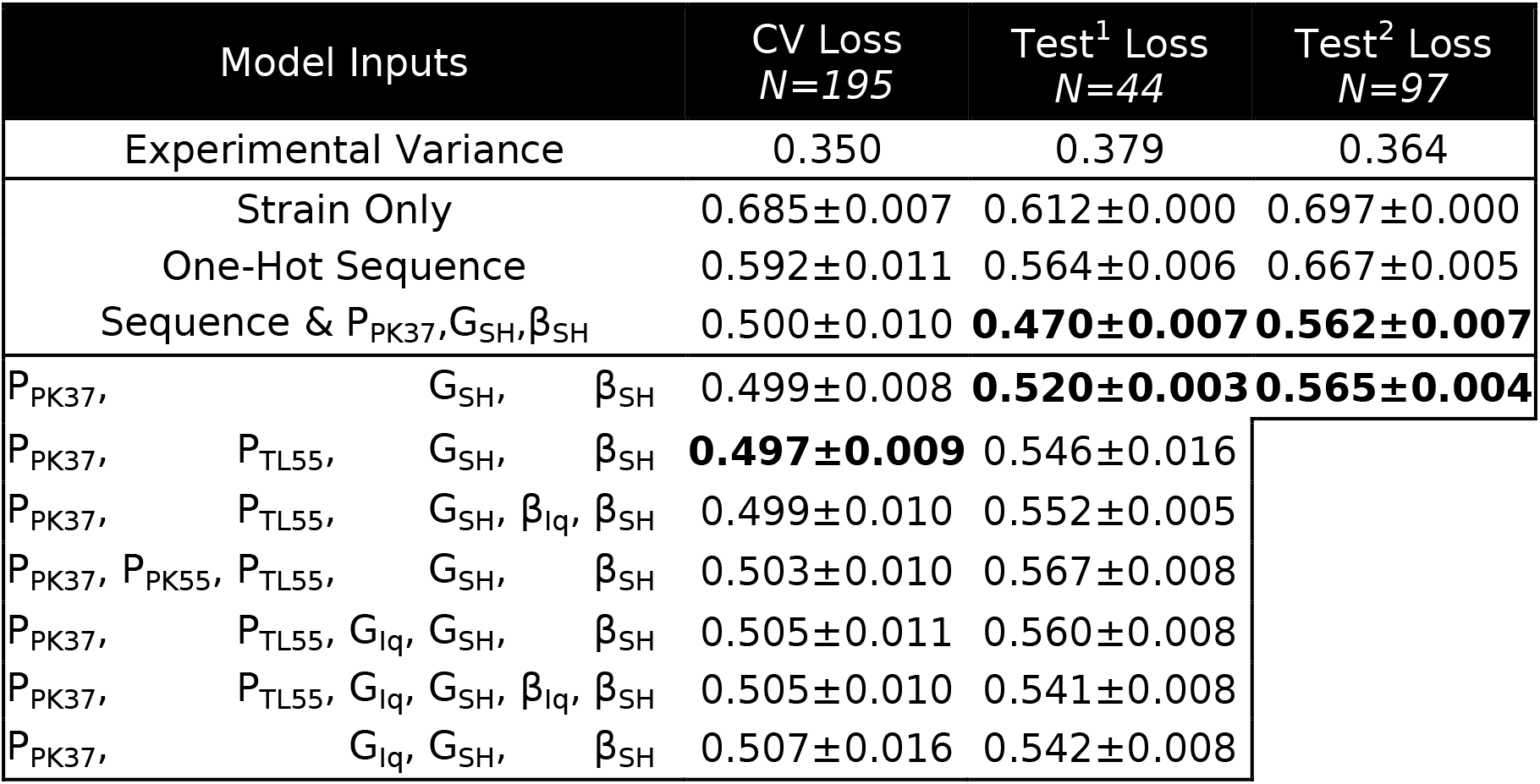
Tabulated performance of models (mean squared error ± standard deviation, *CV*: n=10 repetitions of 10-fold cross-validation. *Test*: n=10 models of same architecture with varying random state during training). CV predicted the yield of 195 unique sequences (I^q^: 73, SH: 122, Both: 39). Test^1^ (44 unique, I^q^: 26, SH 21, Both: 3) and Test^2^ (97 unique, I^q^: 57, SH: 46, Both: 6) sequences were independent of model training and were evaluated to determine the model generalization.

**Supplementary Figure 8.**
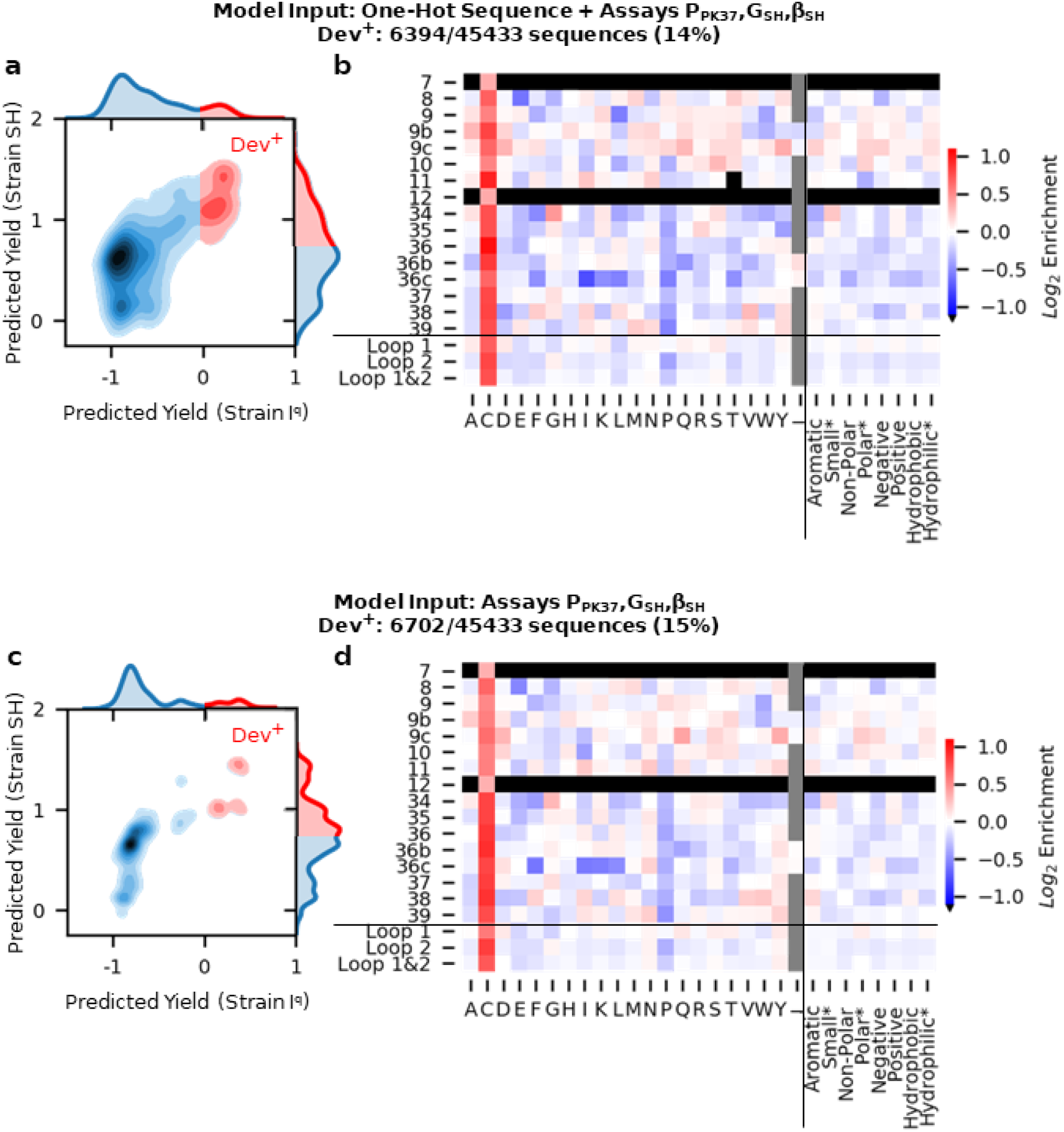
Dev^+^ sitewise amino acid enrichments displays cysteine preference for models when utilizing HT assay scores, with (a,b) and without (c,d) sequence information. The predictive performance of both sets of model inputs was not statistically different. This analysis demonstrates the enrichment of cysteine was trained via HT assays, rather than an effect of the inclusion of sequence information in the model.

**Supplementary Figure 9.**
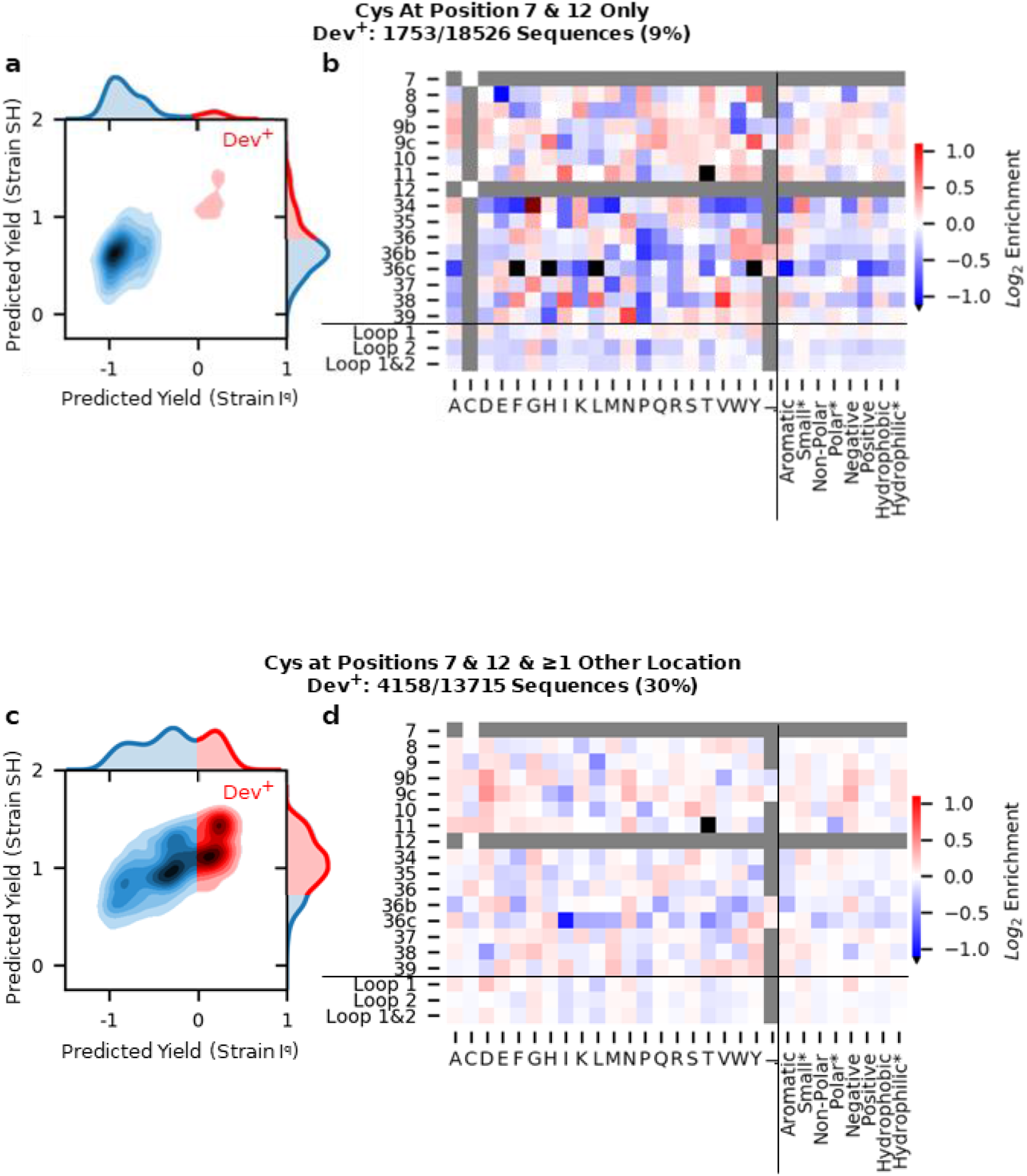
Sitewise enrichment is modified depending on cysteine inclusion outside of positions 7 and 12. Dev^+^ sequences were predicted to have a transformed yield >0 for strain I^q^ and >0.75 for Strain SH. Sitewise amino acid enrichment was calculated as the log2 change in frequency of Dev^+^ versus all predicted sequences. **a,b)** Predicted recombinant yield and enriched amino acids for sequences containing cysteines only at positions 7 and 12. **c,d)** Predicted recombinant yield and enriched amino acids for sequence containing cysteines at positions 7 and 12 and at least one other position.

**DNA Table A:**
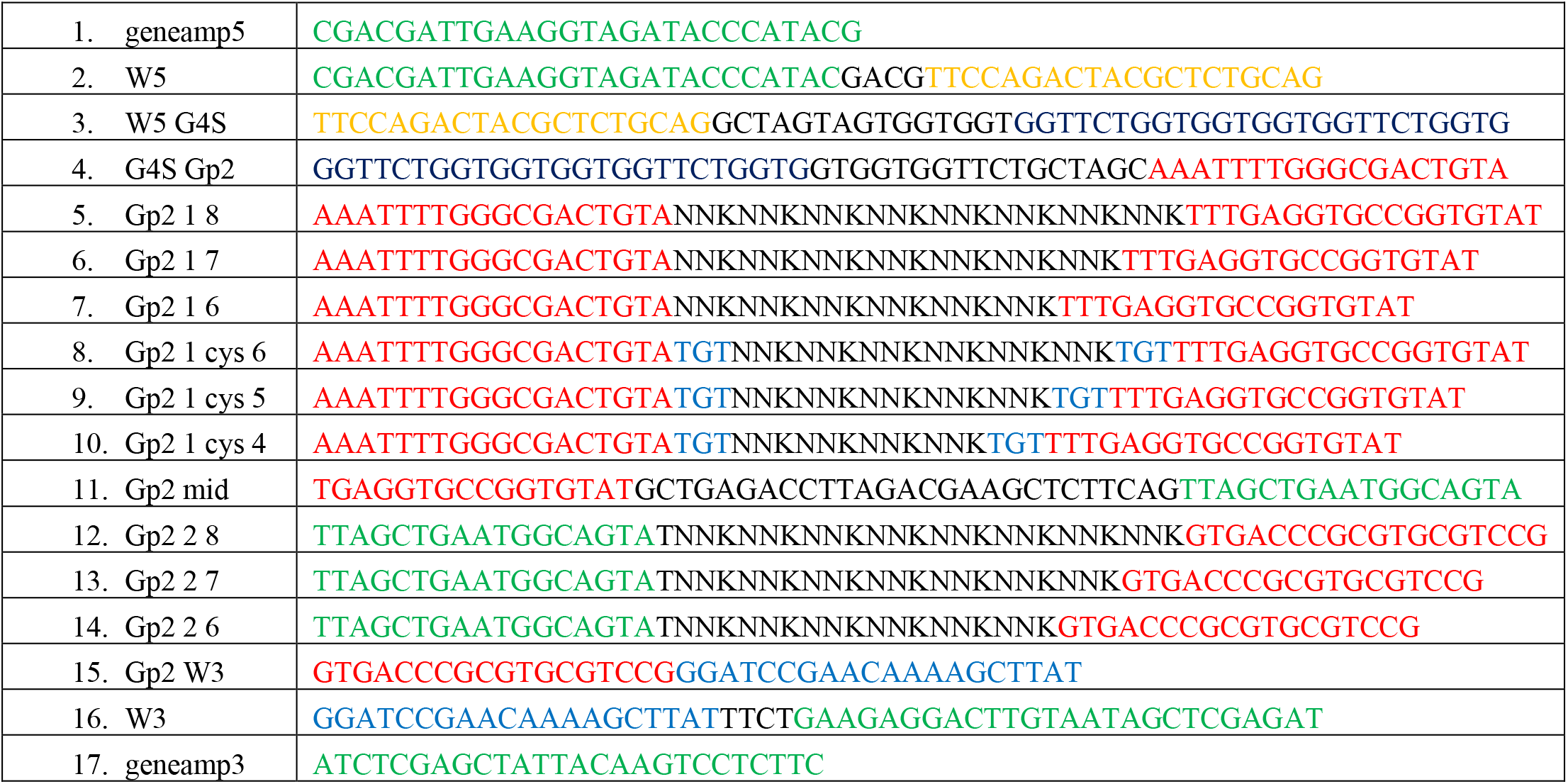
Primers for Gp2 library construction via PCR addition/amplification

**DNA Table B:**
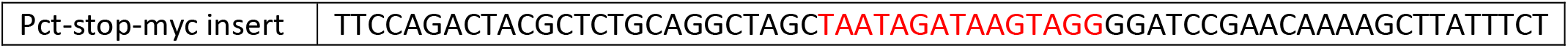
Primer used to create negative control/baseline vector for on-yeast protease screening

**DNA Table C:**
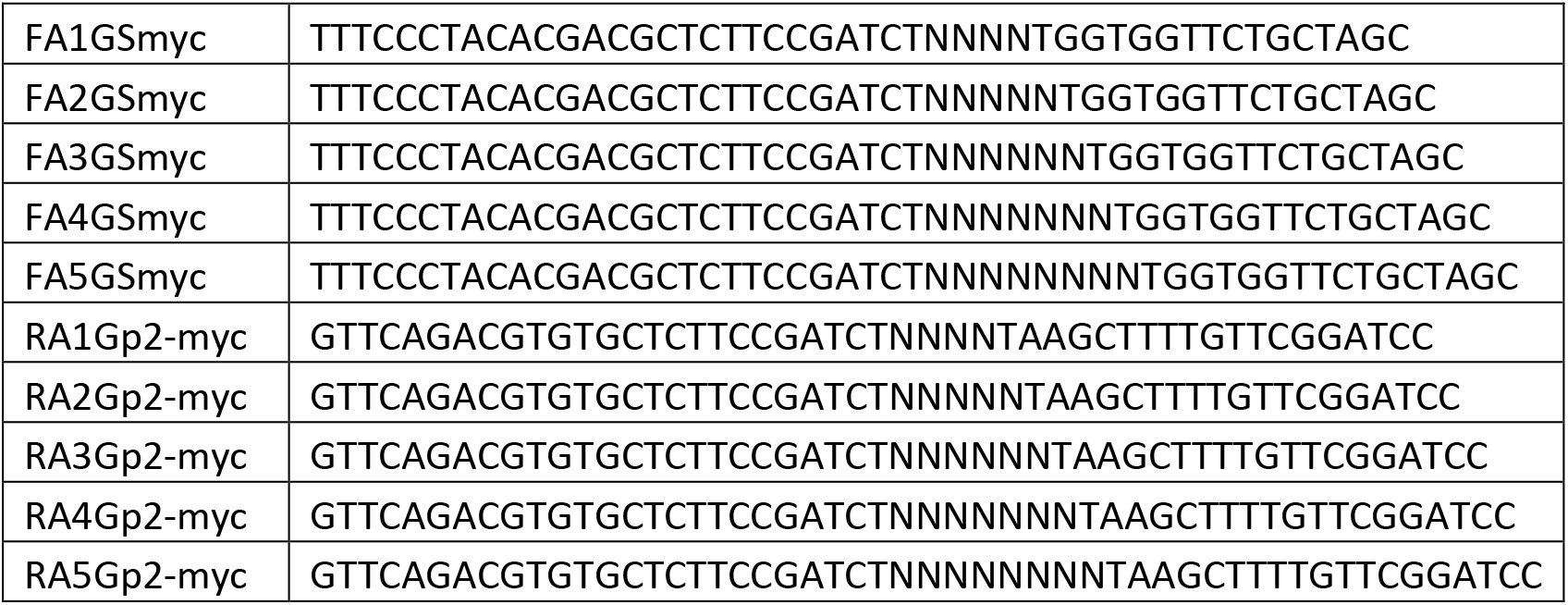
PCR1 primers for illumina preparation of on-yeast protease screening

**DNA Table D:**
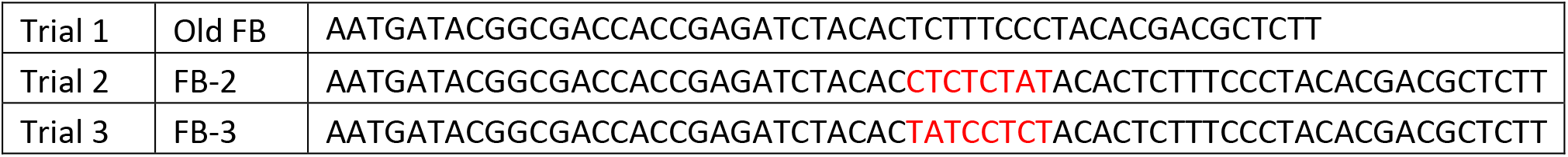
PCR2 Forward primers for illumina sequencing with trial specific barcodes (red). Note Trial 1 was run on a separate Novaseq and thus did not have an FB barcode

**DNA Table E:**
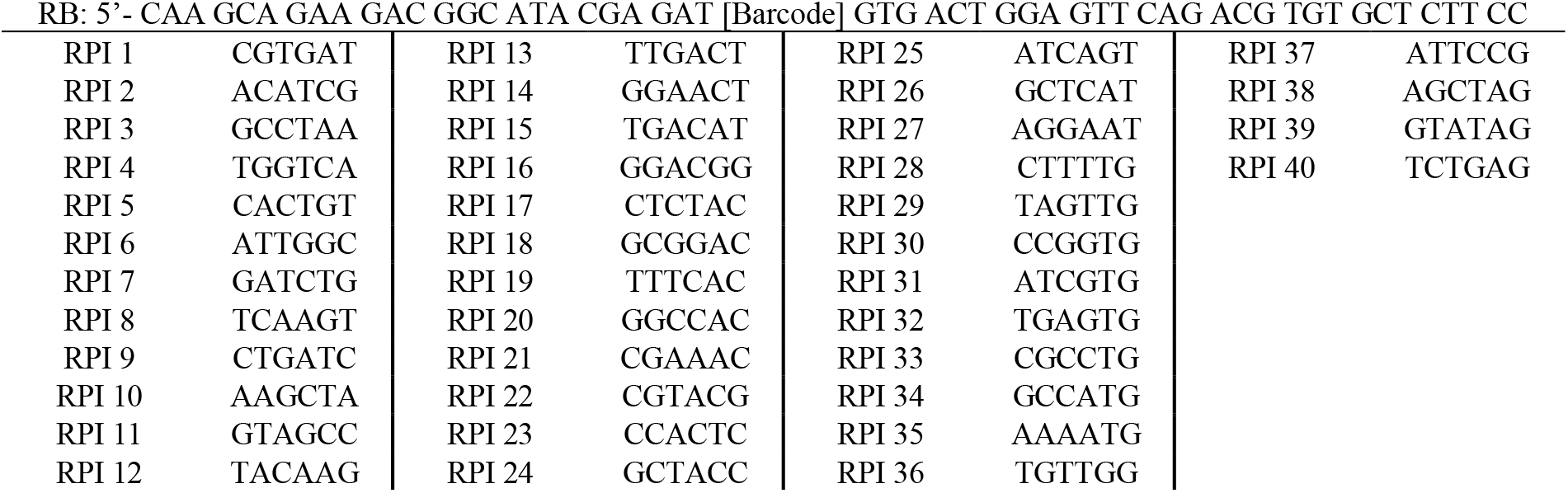
PCR2 reverse primers for illumina sequencing with gate specific barcodes.

**DNA Table F:**
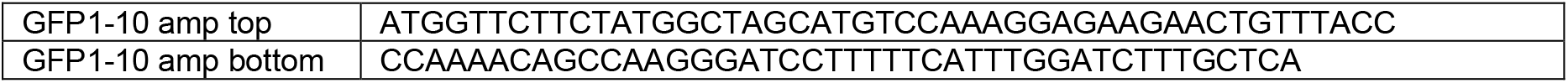
Primers used to amplify GFP1-10 from obtained plamid with overlaps to allow for Gibson assembly into pBAD plasmid

**DNA Table G:**
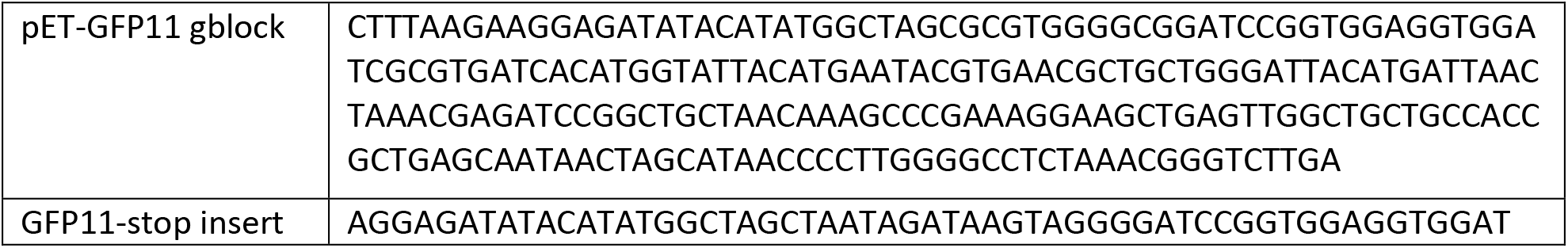
DNA used to add GFP11 to pet (gblock) and to add the stop codon for the negative control

**DNA Table H:**
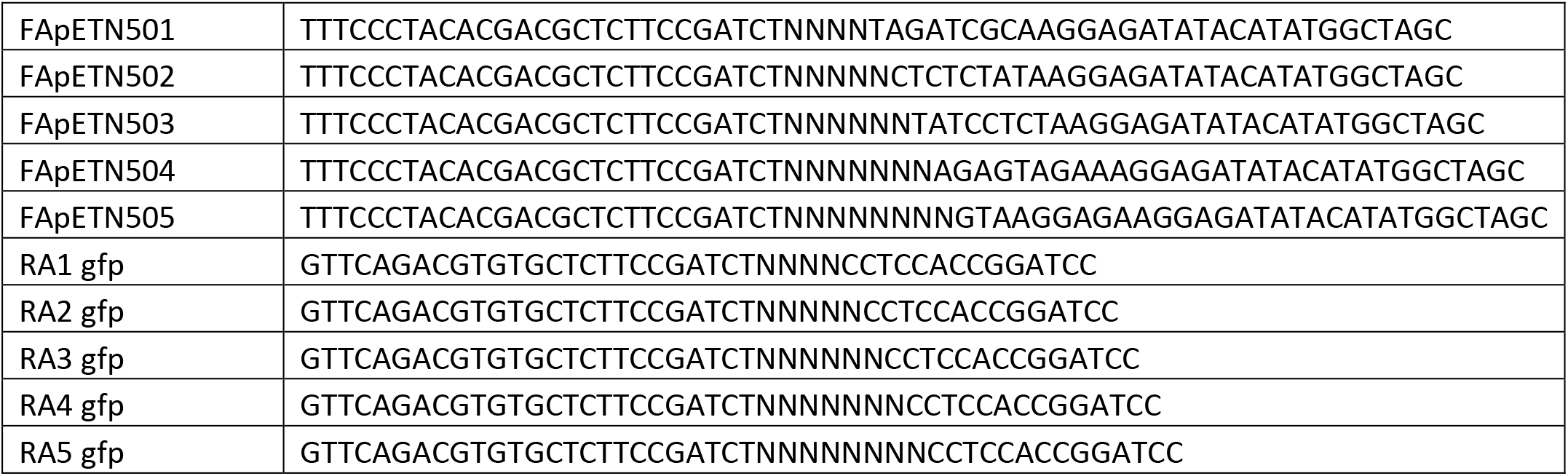
PCR1 primers for illumina preparation of split-GFP assay

**DNA Table I:**
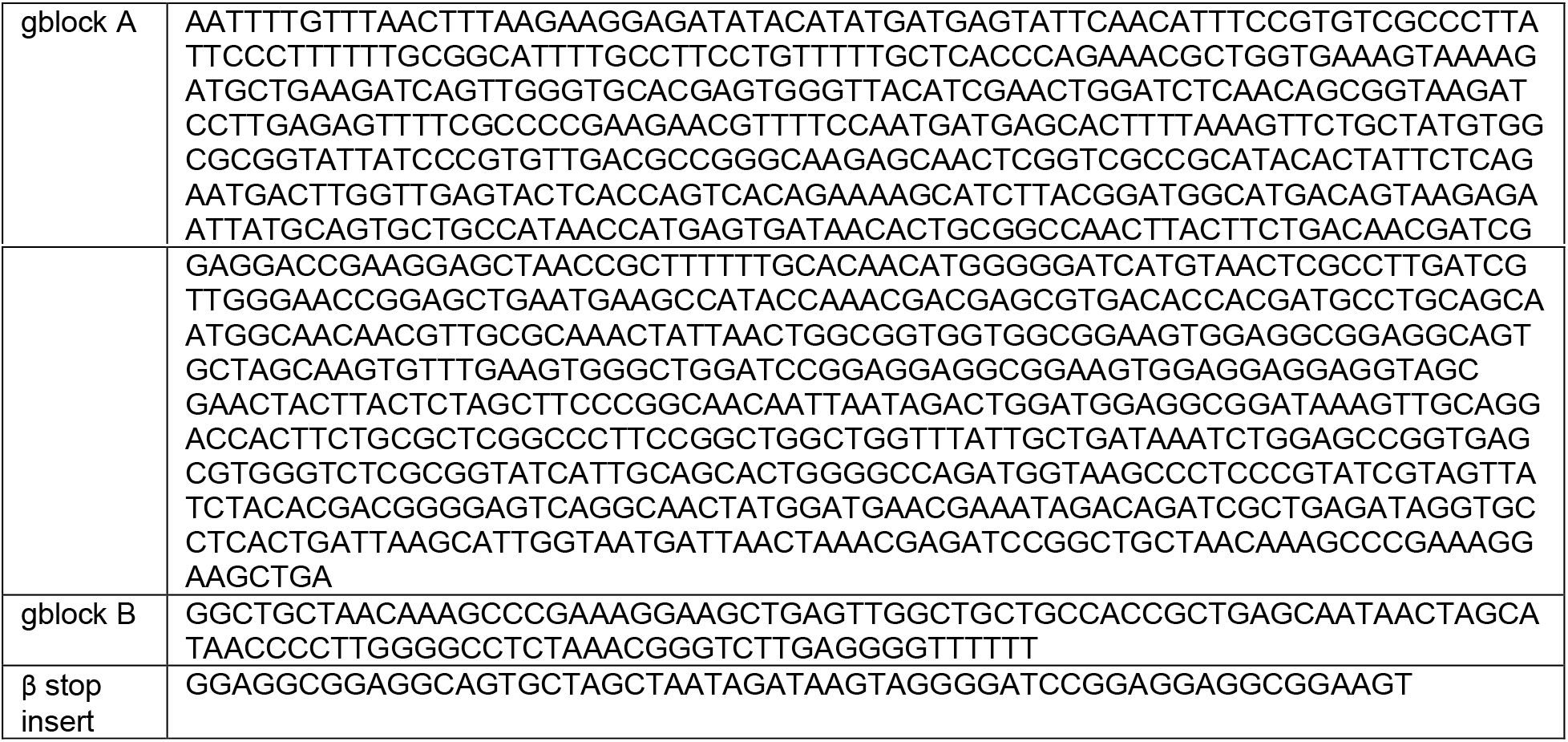
gBlocks for split beta-lactamase plasmid

**DNA Table J:**
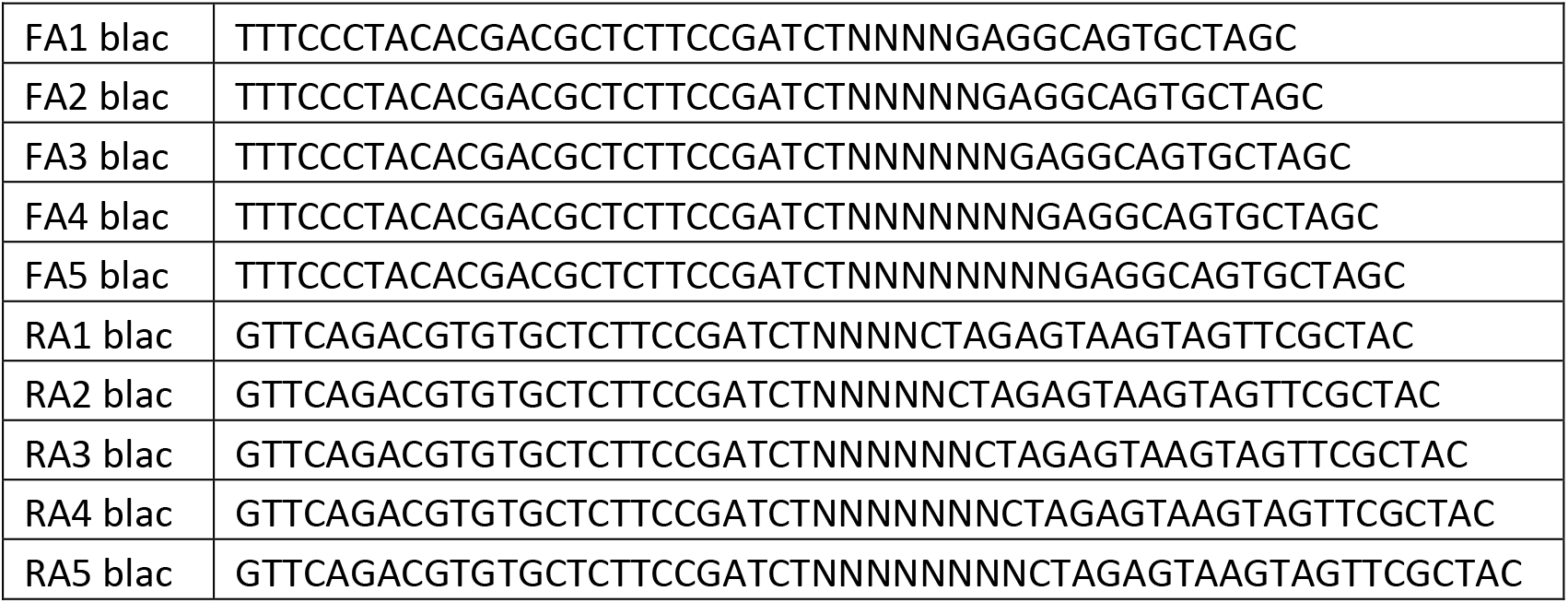
DNA Table J: PCR1 primers for illumina preparation of split-beta lactamase assay

**DNA Table K:**
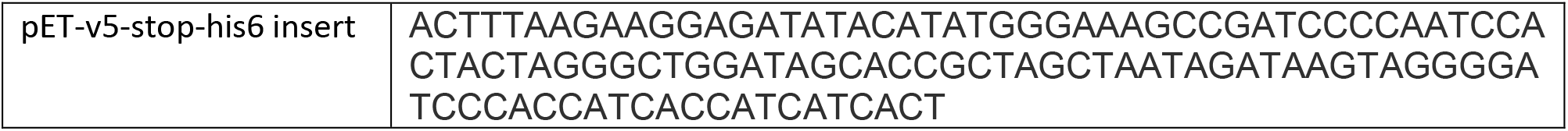
Insert to create pET-V5-stop-His6

**DNA Table L:**
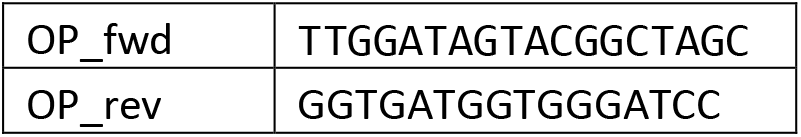
PCR primers to amplify the Twist Oligopool (see Oligopool.fasta)

**DNA Table M:**
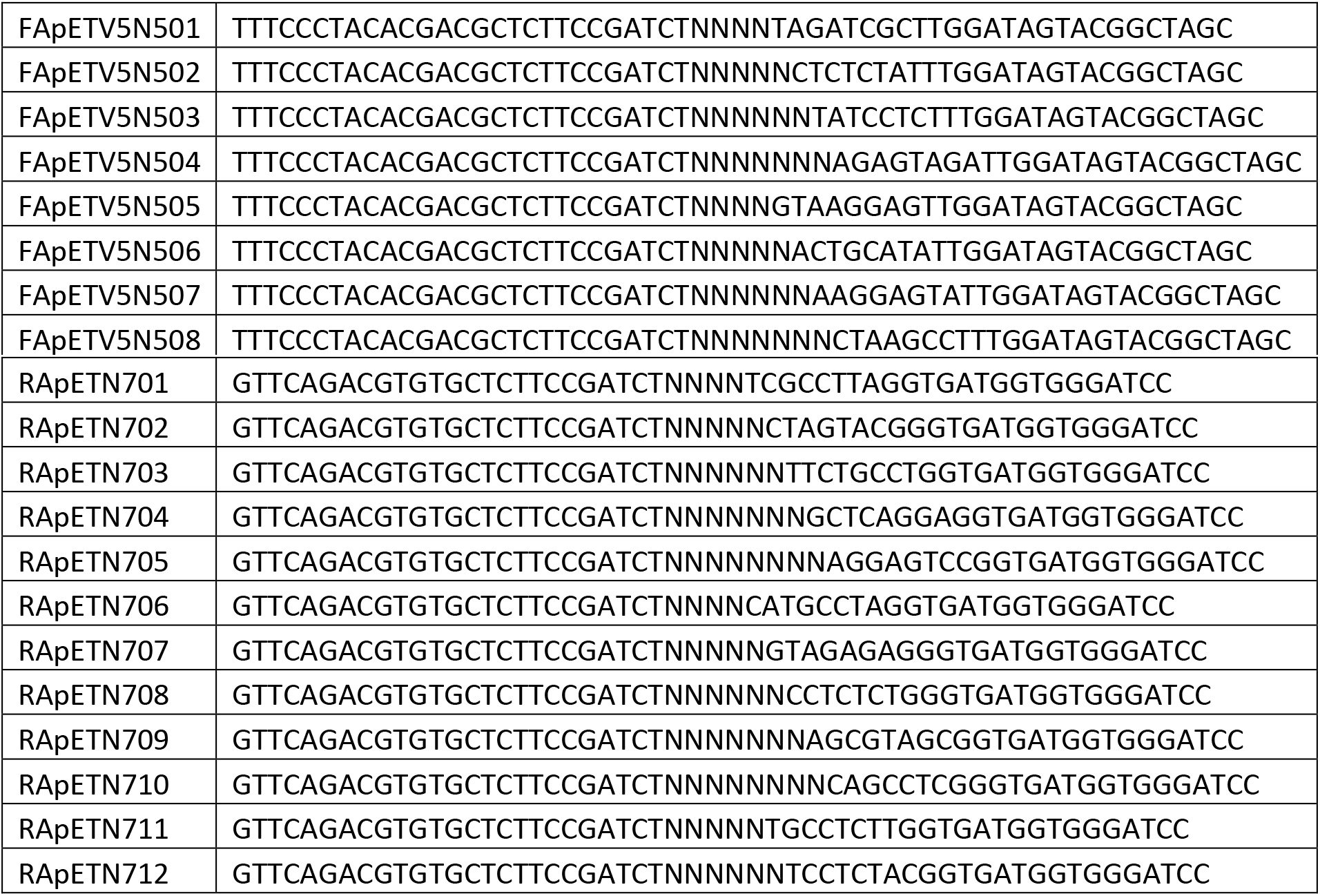
PCR1 primers with row/column specific barcodes to identify the location of the sequence

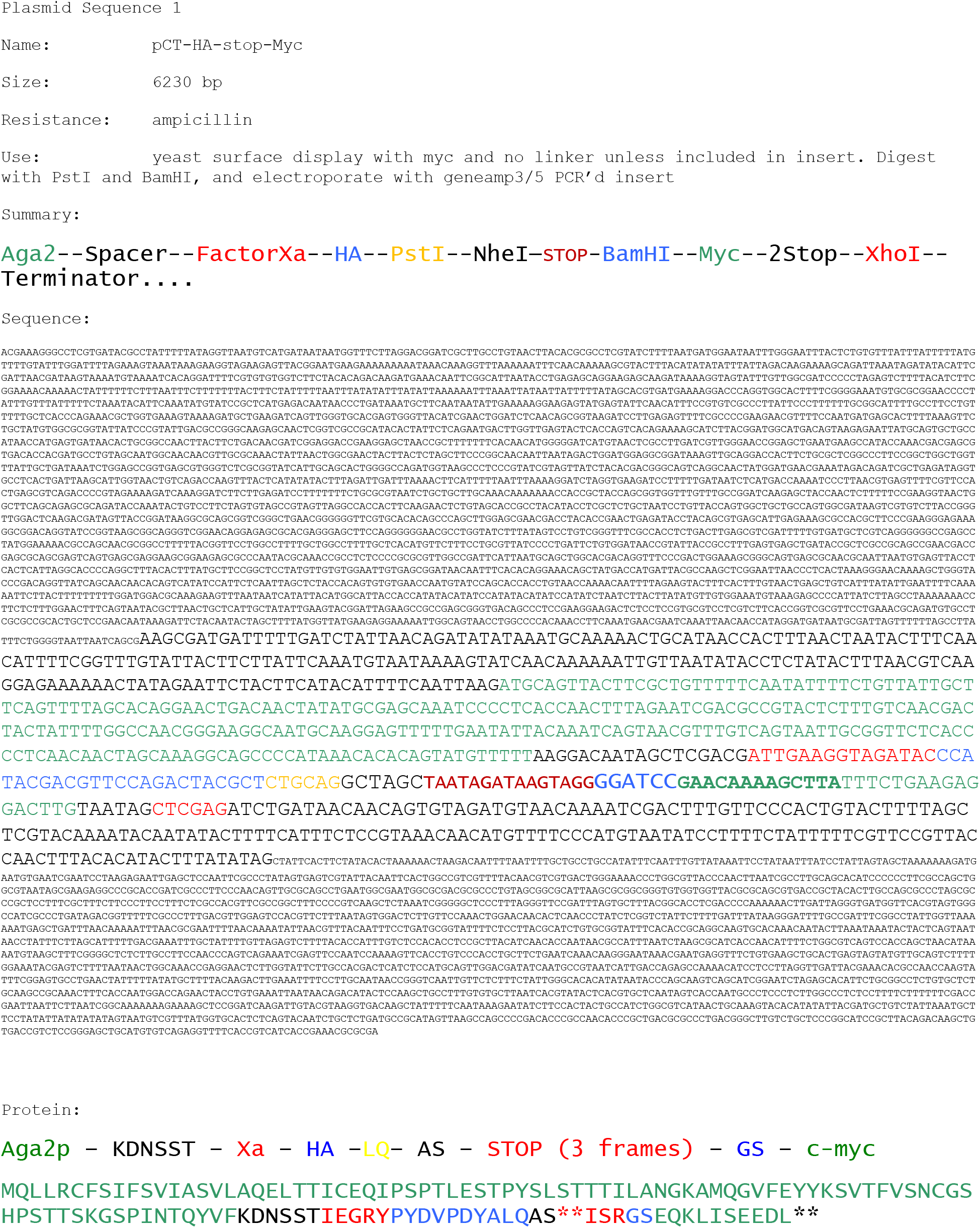

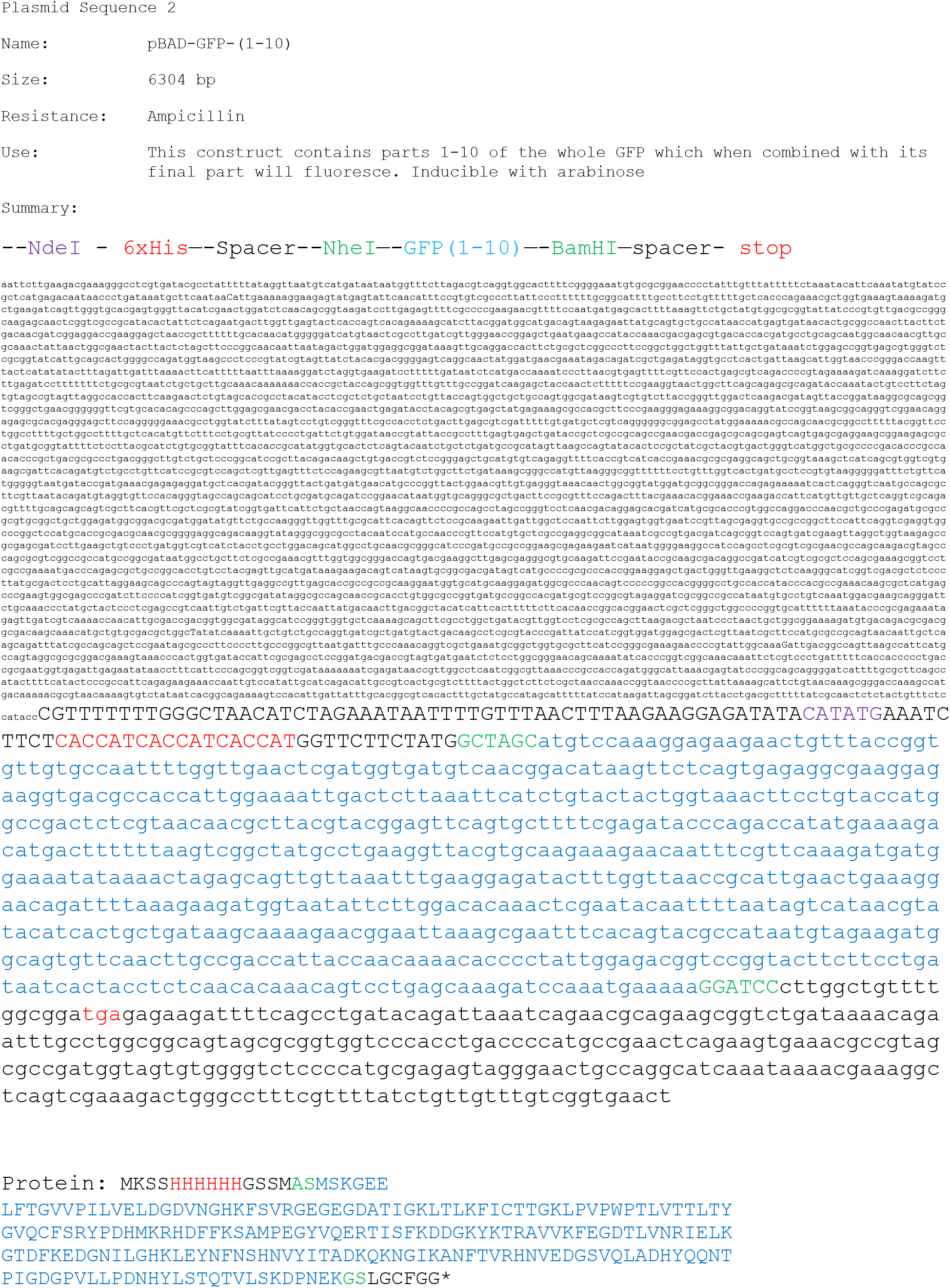

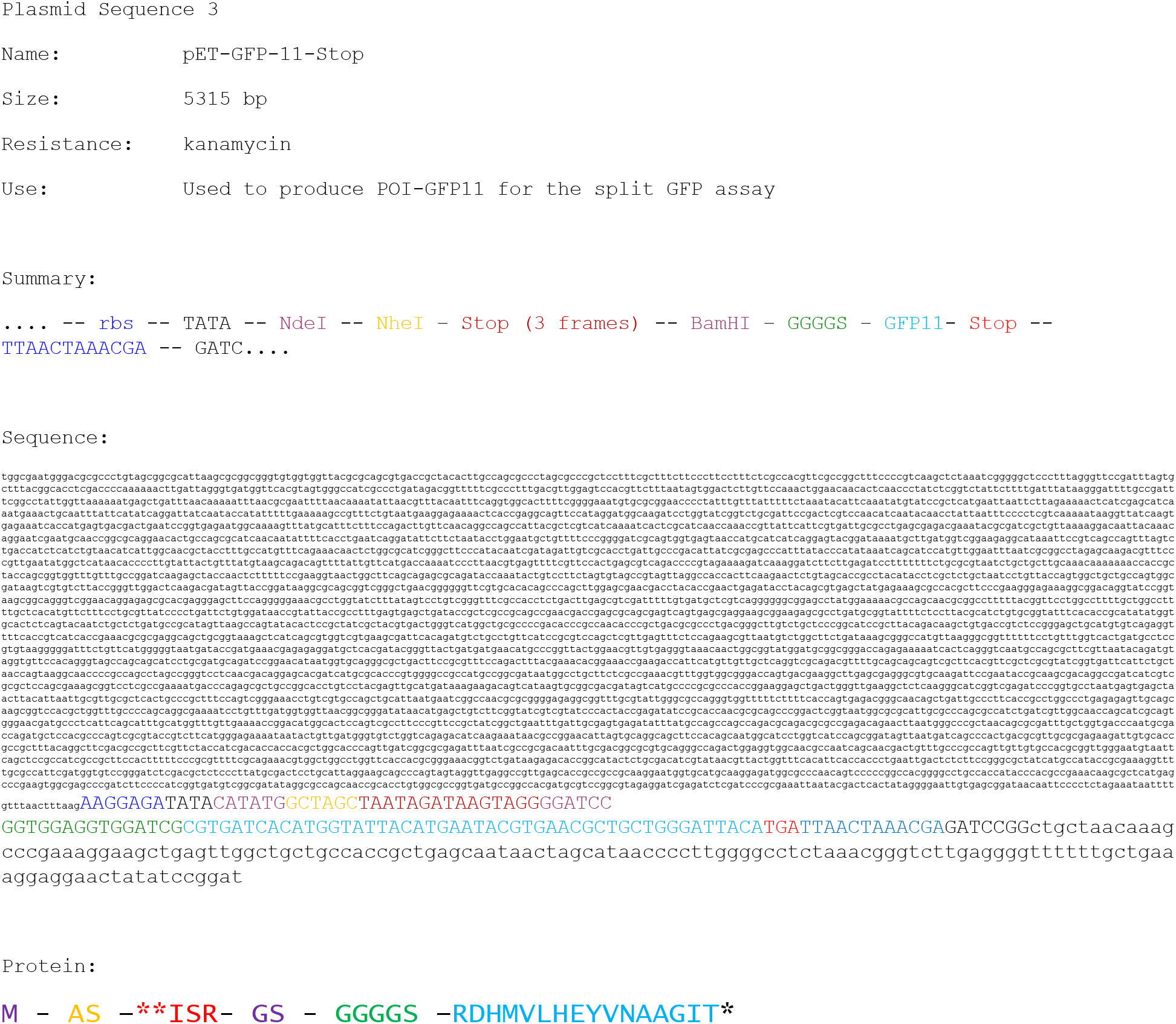

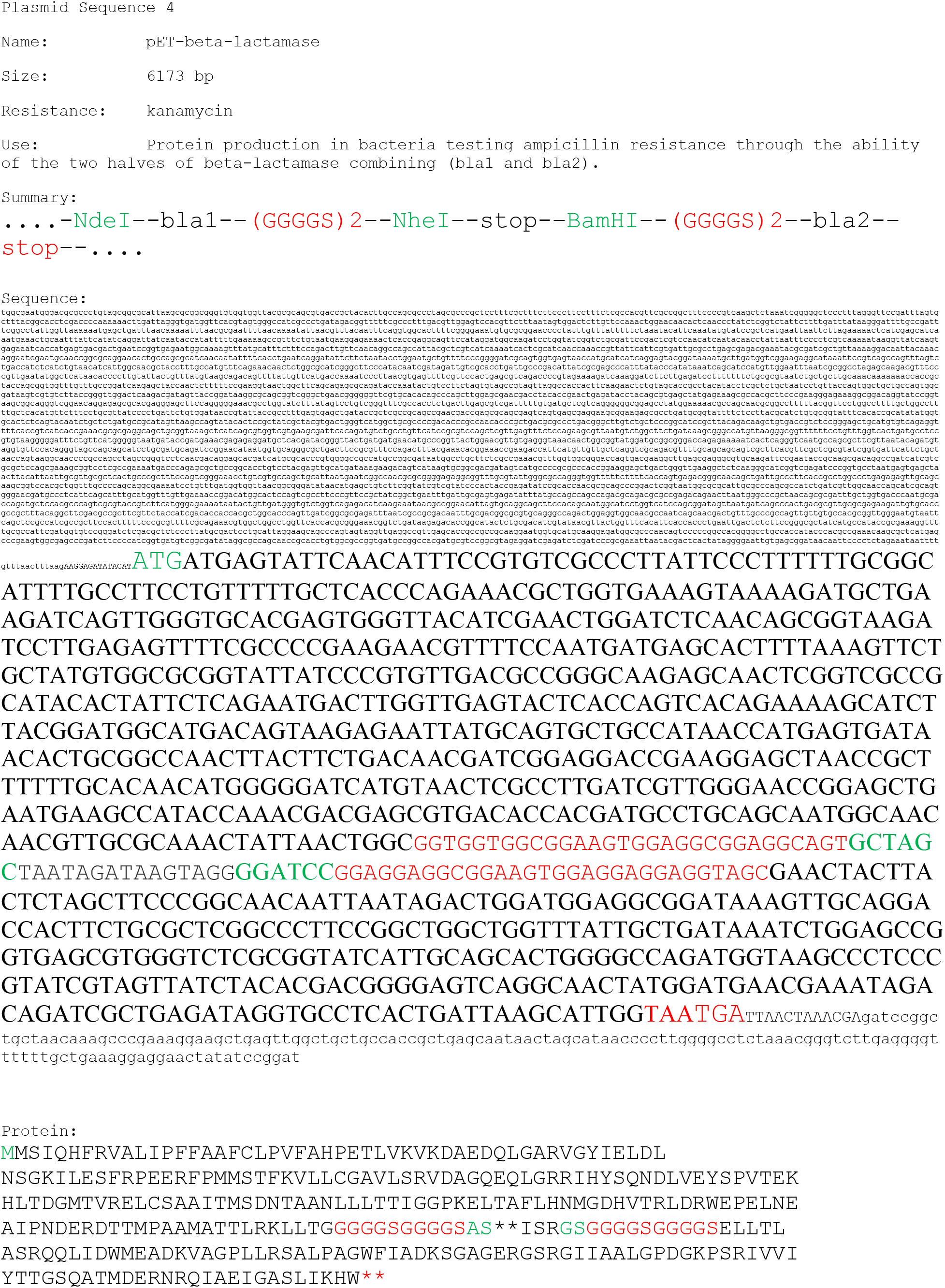

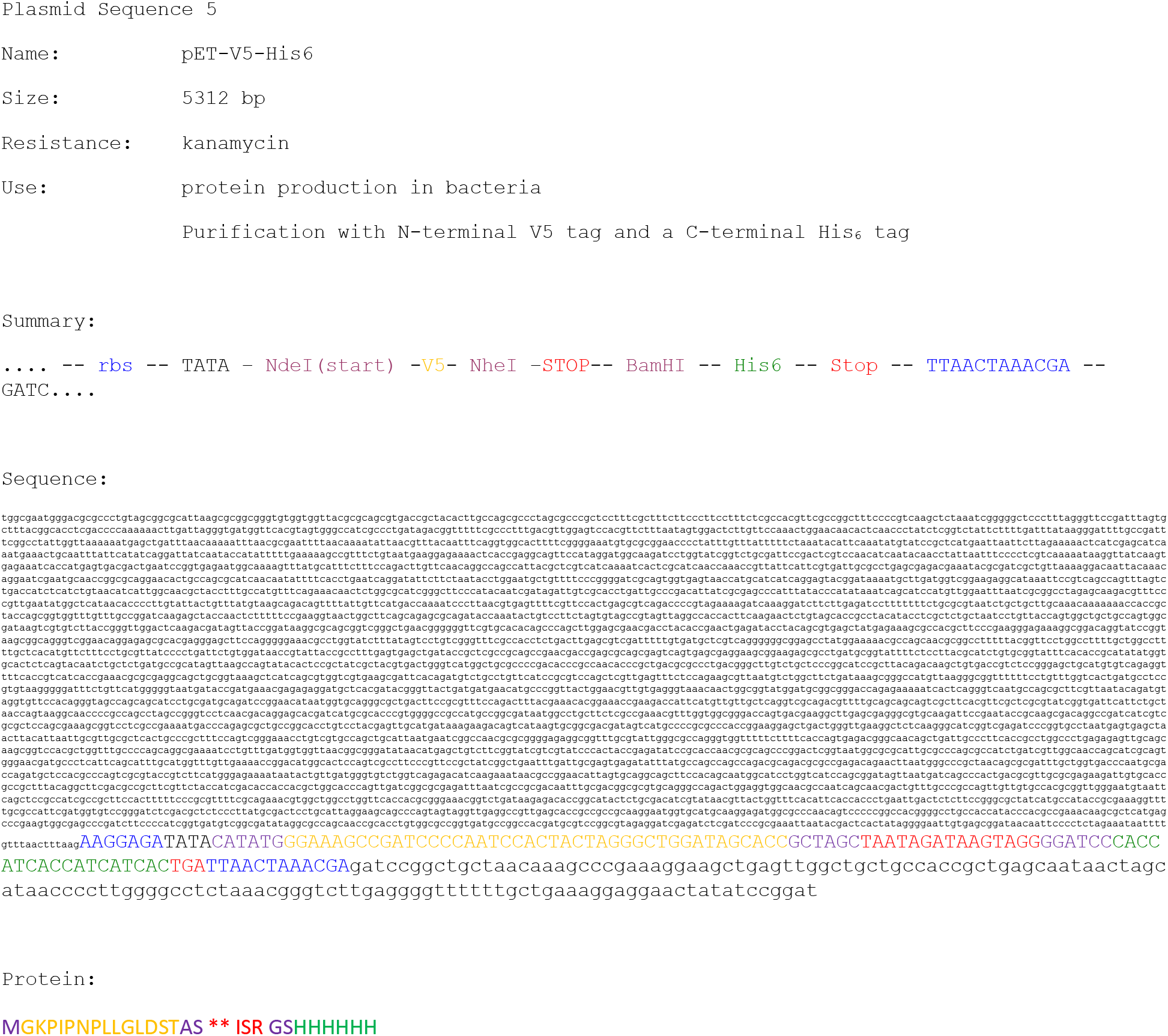

## References

1. Jain, T. et al. Biophysical properties of the clinical-stage antibody landscape. Proc. Natl. Acad. Sci. U. S. A. 114, 944–949 (2017).

2. Xu, Y. et al. Structure, heterogeneity and developability assessment of therapeutic antibodies. MAbs 11, 239–264 (2019).

3. Raybould, M. I. J. et al. Five computational developability guidelines for therapeutic antibody profiling. Proc. Natl. Acad. Sci. U. S. A. 116, 4025–4030 (2019).

4. Yang, X. et al. Developability studies before initiation of process development. MAbs 5, 787–794 (2013).

5. Bailly, M. et al. Predicting Antibody Developability Profiles Through Early Stage Discovery Screening. MAbs 12, (2020).

6. Pérez, A. W. et al. In vitro and in silico assessment of the developability of a designed monoclonal antibody library. MAbs 11, 388–400 (2019).

7. Almagro, J. C., Pedraza-Escalona, M., Arrieta, H. I. & Pérez-Tapia, S. M. Phage Display Libraries for Antibody Therapeutic Discovery and Development. Antibodies (Basel, Switzerland) 8, 44 (2019).

8. Delmar, J. A., Wang, J., Choi, S. W., Martins, J. A. & Mikhail, J. P. Machine Learning Enables Accurate Prediction of Asparagine Deamidation Probability and Rate. Mol. Ther. - Methods Clin. Dev. 15, 264–274 (2019).

9. Lu, X. et al. Deamidation and isomerization liability analysis of 131 clinical-stage antibodies. MAbs 11, 45–57 (2019).

10. Buck, P. M. et al. Computational methods to predict therapeutic protein aggregation. Methods Mol. Biol. 899, 425–451 (2012).

11. Magnan, C. N., Randall, A. & Baldi, P. SOLpro: accurate sequence-based prediction of protein solubility. Bioinformatics 25, 2200–2207 (2009).

12. Agrawal, N. J. et al. Computational tool for the early screening of monoclonal antibodies for their viscosities. MAbs 8, 43–48 (2016).

13. Raybould, M. I. J. et al. Five computational developability guidelines for therapeutic antibody profiling. Proc. Natl. Acad. Sci. 116, 4025 LP–4030 (2019).

14. Chennamsetty, N., Voynov, V., Kayser, V., Helk, B. & Trout, B. L. Design of therapeutic proteins with enhanced stability. Proc. Natl. Acad. Sci. 106, 11937 LP–11942 (2009).

15. Lauer, T. M. et al. Developability index: A rapid in silico tool for the screening of antibody aggregation propensity. J. Pharm. Sci. 101, 102–115 (2012).

16. Potapov, V., Cohen, M. & Schreiber, G. Assessing computational methods for predicting protein stability upon mutation: good on average but not in the details. Protein Eng. Des. Sel. 22, 553–560 (2009).

17. Nisthal, A., Wang, C. Y., Ary, M. L. & Mayo, S. L. Protein stability engineering insights revealed by domain-wide comprehensive mutagenesis. Proc. Natl. Acad. Sci. 116, 16367 LP–16377 (2019).

18. Estep, P. et al. An alternative assay to hydrophobic interaction chromatography for high-throughput characterization of monoclonal antibodies. MAbs 7, 553–561 (2015).

19. Chai, Q. et al. Development of a high-throughput solubility screening assay for use in antibody discovery. MAbs 11, 747–756 (2019).

20. Carter, P. J. & Lazar, G. A. Next generation antibody drugs: pursuit of the ‘ high-hanging fruit ‘. Nat. Rev. Drug Discov. 17, 197–223 (2018).

21. Banta, S., Dooley, K. & Shur, O. Replacing antibodies: engineering new binding proteins. Annu. Rev. Biomed. Eng. 15, 93–113 (2013).

22. Hackel, B. J. Alternative Protein Scaffolds for Molecular Imaging and Therapy. in Engineering in Translational Medicine 343–364 (Springer London, 2014). doi:10.1007/978-1-4471-4372-7_13

23. Stern, L. A., Case, B. A. & Hackel, B. J. Alternative Non-Antibody Protein Scaffolds for Molecular Imaging of Cancer. Curr. Opin. Chem. Eng. 2, 425–432 (2013).

24. Vazquez-Lombardi, R. et al. Challenges and opportunities for non-antibody scaffold drugs. Drug Discov. Today 20, 1271–1283 (2015).

25. Woldring, D. R., Holec, P. V., Stern, L. A., Du, Y. & Hackel, B. J. A Gradient of Sitewise Diversity Promotes Evolutionary Fitness for Binder Discovery in a Three-Helix Bundle Protein Scaffold. Biochemistry 56, 1656–1671 (2017).

26. Miersch, S. & Sidhu, S. S. Synthetic antibodies: Concepts, potential and practical considerations. Methods 57, 486–498 (2012).

27. Kruziki, M. A., Bhatnagar, S., Woldring, D. R., Duong, V. T. & Hackel, B. J. A 45- Amino-Acid Scaffold Mined from the PDB for High-Affinity Ligand Engineering. Chem. Biol. 22, 946–956 (2015).

28. Kruziki, M. A., Sarma, V. & Hackel, B. J. Constrained Combinatorial Libraries of Gp2 Proteins Enhance Discovery of PD-L1 Binders. ACS Comb. Sci. 20, 423–435 (2018).

29. Chan, J. Y., Hackel, B. J. & Yee, D. Targeting Insulin Receptor in Breast Cancer Using Small Engineered Protein Scaffolds. Mol. Cancer Ther. 16, 1324 LP–1334 (2017).

30. Case, B. A., Kruziki, M. A., Johnson, S. M. & Hackel, B. J. Engineered Charge Redistribution of Gp2 Proteins through Guided Diversity for Improved PET Imaging of Epidermal Growth Factor Receptor. Bioconjug. Chem. 29, 1646–1658 (2018).

31. Du, F. et al. Engineering an EGFR-binding Gp2 domain for increased hydrophilicity. Biotechnol. Bioeng. 116, 526–535 (2019).

32. Golinski, A. W., Holec, P. V, Mischler, K. M. & Hackel, B. J. Biophysical Characterization Platform Informs Protein Scaffold Evolvability. ACS Comb. Sci. 21, 323–335 (2019).

33. Rocklin, G. J. et al. Global analysis of protein folding using massively parallel design, synthesis, and testing. Science 357, 168–175 (2017).

34. Ritter, S. C. & Hackel, B. J. Validation and stabilization of a prophage lysin of Clostridium perfringens by using yeast surface display and coevolutionary models. Appl. Environ. Microbiol. 85, (2019).

35. Klesmith, J. R. et al. Retargeting CD19 Chimeric Antigen Receptor T Cells via Engineered CD19-Fusion Proteins. Mol. Pharm. 16, 3544–3558 (2019).

36. Cabantous, S. & Waldo, G. S. In vivo and in vitro protein solubility assays using split GFP. Nat. Methods 3, 845–854 (2006).

37. Foit, L. et al. Optimizing Protein Stability In Vivo. Mol. Cell 36, 861–871 (2009).

38. Overton, T. W. Recombinant protein production in bacterial hosts. Drug Discov. Today 19, 590–601 (2014).

39. Tian, G. et al. Quantitative dot blot analysis (QDB), a versatile high throughput immunoblot method. Oncotarget 8, 58553–58562 (2017).

40. Chen, J. et al. Chaperone activity of DsbC. J. Biol. Chem. 274, 19601–19605 (1999).

41. Boder, E. T. & Wittrup, K. D. Yeast surface display for screening combinatorial polypeptide libraries. Nat. Biotechnol. 15, 553–557 (1997).

42. Galarneau, A., Primeau, M., Trudeau, L.-E. & Michnick, S. W. β-Lactamase protein fragment complementation assays as in vivo and in vitro sensors of protein–protein interactions. Nat. Biotechnol. 20, 619–622 (2002).

43. Mansell, T. J., Linderman, S. W., Fisher, A. C. & DeLisa, M. P. A rapid protein folding assay for the bacterial periplasm. Protein Sci. 19, 1079–1090 (2010).

44. Yeo, I.-K. & Johnson, R. A. A New Family of Power Transformations to Improve Normality or Symmetry. Biometrika 87, 954–959 (2000).

45. Hall, M. A. Correlation-based feature selection for machine learning. (1999).

46. Hoffmann, F. & Rinas, U. Stress Induced by Recombinant Protein Production in Escherichia coli BT - Physiological Stress Responses in Bioprocesses: -/-. in 73–92 (Springer Berlin Heidelberg, 2004). doi:10.1007/b93994

47. Hackel, B. J., Ackerman, M. E., Howland, S. W. & Wittrup, K. D. Stability and CDR Composition Biases Enrich Binder Functionality Landscapes. J. Mol. Biol. 401, 84–96 (2010).

48. Fellouse, F. A. et al. High-throughput generation of synthetic antibodies from highly functional minimalist phage-displayed libraries. J. Mol. Biol. 373, 924–940 (2007).

49. Seeger, M. A. et al. Design, construction, and characterization of a second-generation DARPin library with reduced hydrophobicity. Protein Sci. 22, 1239–1257 (2013).

50. Koide, A., Wojcik, J., Gilbreth, R. N., Hoey, R. J. & Koide, S. Teaching an Old Scaffold New Tricks: Monobodies Constructed Using Alternative Surfaces of the FN3 Scaffold. J. Mol. Biol. 415, 393–405 (2012).

## Methods References

1. Kruziki, M. A., Sarma, V. & Hackel, B. J. Constrained Combinatorial Libraries of Gp2 Proteins Enhance Discovery of PD-L1 Binders. ACS Comb. Sci. 20, 423–435 (2018).

2. Chao, G. et al. Isolating and engineering human antibodies using yeast surface display. Nat. Protoc. 1, 755–768 (2006).

3. Scanlon, T. C., Gray, E. C. & Griswold, K. E. Quantifying and resolving multiple vector transformants in S. cerevisiae plasmid libraries. BMC Biotechnol. 9, 95 (2009).

4. Kruziki, M. A., Bhatnagar, S., Woldring, D. R., Duong, V. T. & Hackel, B. J. A 45- Amino-Acid Scaffold Mined from the PDB for High-Affinity Ligand Engineering. Chem. Biol. 22, 946–956 (2015).

5. Kamiyama, D. et al. Versatile protein tagging in cells with split fluorescent protein. Nat. Commun. 7, 11046 (2016).

6. Golinski, A. W., Holec, P. V, Mischler, K. M. & Hackel, B. J. Biophysical Characterization Platform Informs Protein Scaffold Evolvability. ACS Comb. Sci. 21, 323–335 (2019).

7. Edgar, R. C. Search and clustering orders of magnitude faster than BLAST. Bioinformatics 26, 2460–2461 (2010).

8. Edgar, R. C. UNOISE2: improved error-correction for Illumina 16S and ITS amplicon sequencing. bioRxiv 81257 (2016). doi:10.1101/081257

9. Schindelin, J. et al. Fiji: an open-source platform for biological-image analysis. Nat. Methods 9, 676–682 (2012).

10. Schneider, C. A., Rasband, W. S. & Eliceiri, K. W. NIH Image to ImageJ: 25 years of image analysis. Nat. Methods 9, 671–675 (2012).

11. Bergstra, J., Yamins, D. & Cox, D. D. Making a Science of Model Search: Hyperparameter Optimization in Hundreds of Dimensions for Vision Architectures. in Proceedings of the 30th International Conference on International Conference on Machine Learning - Volume 28 I–115–I–123 (JMLR.org, 2013).

12. Bergstra, J., Komer, B., Eliasmith, C., Yamins, D. & Cox, D. D. Hyperopt: a python library for model selection and hyperparameter optimization. Comput. Sci. Discov. 8, 14008 (2015).

13. Hall, M. A. Correlation-based feature selection for machine learning. (1999).

14. Lebigot, E. O. Uncertainties: a Python package for calculations with uncertainties. URL http://pythonhosted.org/uncertainties (2010).

